# Genome-wide analysis of DNA uptake across the outer membrane of naturally competent *Haemophilus influenzae*

**DOI:** 10.1101/866426

**Authors:** Marcelo Mora, Joshua Chang Mell, Garth D. Ehrlich, Rachel L. Ehrlich, Rosemary J. Redfield

**Affiliations:** Department of Zoology, University of British Columbia, Vancouver, BC, Canada V6T 1Z3.; Department of Microbiology & Immunology; Center for Genomic Sciences, Institute of Molecular Medicine and Infectious Disease; Drexel University College of Medicine, 12 Philadelphia PA, USA 19102.; Department of Otolaryngology – Head and Neck Surgery, Drexel University College of Medicine, 12 Philadelphia PA, USA 19102.

## Abstract

The genomes of naturally competent Pasteurellaceae and Neisseriaceae have many short uptake sequences (USS), which allow them to distinguish self-DNA from foreign DNA. To fully characterize this preference we developed genome-wide maps of DNA uptake using both a sequence-based computational model and genomic DNA that had been sequenced after uptake by and recovery from competent *Haemophilus influenzae* cells. When DNA fragments were shorter than the average USS spacing of ~1000 bp, sharp peaks of uptake were centered at USS and separated by valleys with 1000-fold lower uptake. Long DNA fragments (1.5-17 kb) gave much less variation, with 90% of positions having uptake within two-fold of the mean. All detectable uptake biases arose from sequences that fit the USS uptake motif. Simulated competition predicted that, in its respiratory tract environment, *H. influenzae* will efficiently take up its own DNA even when human DNA is present in 100-fold excess.

## INTRODUCTION

Many bacteria are naturally competent, able to actively bind DNA fragments at the cell surface and pull them into the cytoplasm, where the incoming fragments may contribute nucleotides to cellular pools or recombine with homologous genomic sequences (Lorenz and Wackernagel, 1994). The genetic exchange associated with this latter process contributes to adaptation and is known to have promoted resistance to antibiotics (Bae et al., 2014) and increased strains’ intracellular invasiveness (Mell et al., 2016) and vaccine resistance (Kress-Bennett et al., 2016; Straume et al., 2015). Thus, understanding how different genomic regions evolve via natural transformation processes could be used to predict the spread of pathogenic traits.

Most naturally competent bacteria that have been tested take up DNA regardless of sequence, but species in two families, the Pasteurellaceae and the Neisseriaceae, exhibit strong preferences for DNA containing short sequence motifs (Chen and Dubnau, 2004). Because these motifs have become highly enriched in the corresponding genomes, these biases effectively limit uptake to DNA from close relatives with the same uptake specificity (Dougherty et al., 1979; Scocca et al., 1974). The distribution of the preferred sequences around the chromosome is uneven (Smith et al., 1995), which may cause different genes to experience quite different rates of genetic exchange.

Most steps in the natural transformation process are highly conserved among transformable species (Chen and Dubnau, 2004). In the Pasteurellaceae, the Neisseriaceae and most other Gram-negative bacteria, DNA uptake is initiated by binding of a type IV pilus uptake machine to double-stranded DNA (dsDNA) at the cell surface. The DNA-binding protein has not been identified in *H. influenzae*, but in *Neisseria* it is a minor pilin-type protein that forms part of the pilus (Cehovin et al 2013). DNA binding is followed by retraction of the pilus, which pulls the DNA across the outer membrane into the periplasm. Because circular DNA is taken up as efficiently as linear DNA, uptake is thought to begin internally on DNA fragments rather than at their ends (Barany et al., 1983). Thus, it is likely that the stiff dsDNA molecule is transiently kinked (folded sharply back on itself) at the site of initiation to allow it to pass through the narrow secretin pore of the uptake machinery. Forces generated by the retraction of the type IV pilus are thought to be responsible for this kinking, which might be facilitated by strand separation at the AT-tracts (Danner et al., 1982). Once a loop of the DNA is inside the periplasm, a ratchet process controlled by the periplasmic protein ComEA is thought to pull the rest of the DNA through the outer membrane (Hepp and Maier, 2016; Salzer et al., 2016). Below we use ‘DNA uptake’ to refer to the combined binding and membrane-transport steps that move DNA from the extracellular environment across the outer membrane into the periplasm. In Gram-positive bacteria similar machinery acts to pull the DNA through the thick cell wall (Chen and Dubnau, 2004). The next step in natural transformation is translocation of the DNA out of the periplasm and into the cytoplasm. Only the 3’-leading strand remains intact, passing through an inner membrane pore encoded by the *rec2*/*comEC* gene, while the other strand is degraded in the periplasm and its nucleotides are dephosphorylated and imported as nucleosides (Pifer and Smith, 1985). Circular DNA molecules are efficiently taken up across the outer membrane but remain the periplasm because they lack free ends (Pifer and Smith, 1985). As the single strand enters the cytoplasm it undergoes limited exonucleolytic degradation before being complexed with cellular proteins. If sequence similarity permits the strand may then recombine with homologous chromosomal sequences; otherwise the strand is degraded to its constituent nucleotides (de Vries et al., 2001). Both linkage and sequencing studies indicate that fragments much longer than the cell are readily taken up (Goodgal, 1982; Mell and Redfield, 2014), as are fragments as short as 200 bp, although a lower limit has not been established (Mell 2012, Maughan 2009). However, recombination of short fragments is limited because they are usually degraded by cytoplasmic nucleases before they can recombine (Pifer and Smith, 1985). Uptake speed has been estimated at 500-1000 bp/sec, with transformation essentially complete by 15 min (Deich and Smith 1980).

### Direct measures of DNA uptake bias

Uptake-competition experiments in the Pasteurellacean *Haemophilus influenzae* and in *Neisseria gonorrhoeae* showed that genetically marked ‘self-derived’ DNA competes for uptake with unmarked self-derived DNA but not with DNA from unrelated sources (Dougherty et al., 1979; Scocca et al., 1974). Subsequent DNA uptake experiments using cloned radiolabeled DNA fragments found that these self-preferences are caused by the uptake machineries’ strong biases for short sequence motifs, called uptake signal sequences (USS) in *H. influenzae* and DNA uptake sequences (DUS) in *Neisseria* species (Sisco and Smith, 1979) (Davidsen et al., 2004). Sequence comparisons and site-directed mutagenesis initially identified the *H. influenzae* motif as an 11 bp sequence with a strong contribution by flanking AT-rich sequences (Danner et al., 1980, 1982), and later genome sequencing identified 1465 occurrences of a 9 bp USS core in *H. influenzae* and 1892 occurrences of an unrelated 10 bp DUS motif in *N. meningitidis* (Smith et al., 1995, 1999). Subsequent motif search analyses by Maughan *et al*. (Maughan et al., 2010) expanded this number to 2206 USSs in the genome of the standard *H. influenzae* lab strain Rd, sharing the motif shown in Figure 1A. These genomic analyses were later complemented by direct uptake experiments using mutated and degenerate USS variants(Maughan, Wilson and Redfield, 2010; Mell, Hall and Redfield, 2012).. Mell *et al*. used mutagenesis and sequencing of pools of degenerate USS-containing fragments that had been recovered after uptake to identify the contribution to uptake of each USS position, and found that the central GCGG bases are crucial for uptake, with much smaller contributions made by the flanking bases and AT-rich segments. The motif in Figure 1B shows the contribution of each base considered independently. Interaction effects between bases of the AT-tracts and the core were also found to make important contributions to uptake, but only a few of these have been directly measured. Although characterization of the unrelated Neisseriacean DUS has not reached this level of detail (Mathis and Scocca, 1982), the two uptake systems shares many features, apparently convergently evolved, including the presence within each family of lineages with slightly different preferred motifs (Frye et al., 2013; Redfield et al., 2006).

**Figure 1.**
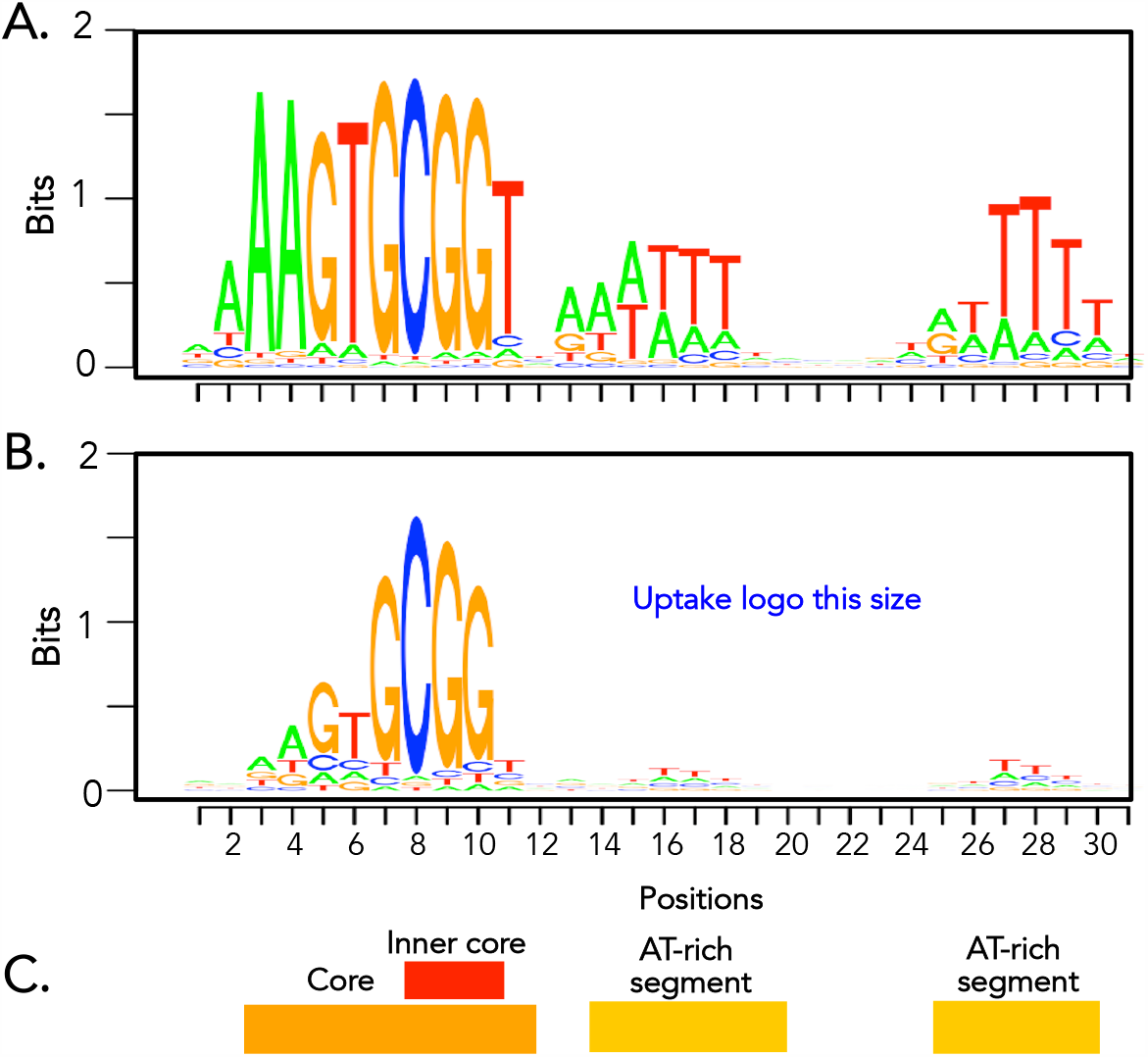
The *H. influenzae* uptake signal sequence. **Legend: A**. Sequence logo showing the individual contributions to genomic abundance of bases in the USS motif (Maughan 2010). **B**. Sequence logo showing the individual contributions to uptake of bases in the USS motif, as measured by Mell *et al*. **C**. Conserved USS segments. See also Figure S1.

### Evolution of uptake sequences in the genome

Alignment of homologous genomic regions from different Pasteurellaceae species showed that USS evolve by point mutations (Redfield et al., 2006); *i*.*e*. they are not inserted elements. As a mechanism for this evolution, (Danner et al., 1980) proposed that the combination of uptake bias and genomic recombination creates an evolutionary pressure that causes the preferred uptake sequences to accumulate throughout the genome, with their numbers limited by their eventual interference with gene function. Consistent with this, uptake sequences in both *Haemophilus* and *Neisseria* are underrepresented in newly acquired segments, in rRNA genes, and in coding sequences, especially those with strong functional constraints (Findlay and Redfield, 2009; Smith et al., 1999). Modeling by Maughan *et al*. (Maughan et al., 2010) confirmed that this molecular drive process could produce uptake sequence distributions like those of real genomes, with no need for any fitness benefit from either the uptake sequences or the recombination they promote. Thus, the presence of biased DNA uptake machinery may be sufficient in itself to explain the abundance of uptake sequences. Such biases may be solely a consequence of direct selection on the DNA uptake machinery for more effective DNA binding or may have been reinforced by indirect selection for preferential uptake of conspecific DNA. These processes might be especially important in respiratory tracts and other mucosal environments where Pasteurellaceae and Neisseriaceae species mainly occur (Man et al., 2017). These environments contain abundant host DNA, and transformation can only occur if the released bacterial DNA competes successfully for binding to the uptake machinery (Lethem et al., 1990; Shak et al., 1990).

The goal of the present study was to measure DNA uptake at every position in the *H. influenzae* genome and to use this uptake data to characterize the effects of USS and identify any other factors affecting uptake. To prepare a framework for interpreting uptake biases, we developed a computational model that predicted the effect of uptake sequences on DNA uptake across the *H. influenzae* genome. We did not attempt to build a model that accurately simulated the actual events of DNA uptake, since too little is known about these. Instead, the initial version of this model was ‘naïve’ in that its parameters and settings were based only on previously published information. Discrepancies between the model’s predictions and the observed uptake were then used to identify features of uptake that were poorly predicted. Hypothesized biological explanations for these discrepancies then guided changes to the model, and the effect of each change on the discrepancy was used to confirm or refute the hypothesis. The most important product of this recursive analysis was not the model itself, but the improved understanding of factors affecting DNA uptake across the genome. These in turn increased the understanding of the genomic distribution of recombination and the effects of competition with DNA from the host or other microbiota.

## RESULTS

### A computational model of DNA uptake

As a framework for interpreting DNA uptake data we developed a simulation model of USS-dependent DNA uptake. It takes as input the locations and strengths of USSs in the DNA whose uptake is to be simulated, the fragment-size distribution of this DNA, and binding and uptake functions that specify how uptake probability depends on USS presence and strength. The output is the expected relative uptake of every position in the genome. Development of the model was guided by basic principles of sequence-specific protein-DNA interactions (Halford and Marko, 2004; Rohs et al., 2010). The first step in these interactions is thought to be a random encounter between a DNA fragment and the binding site of the protein, usually at a DNA position that does not contain the protein’s preferred sequence. This non-specific binding dramatically increases the probability that the protein will subsequently encounter any preferred sequence, either by sliding along the DNA or by transient dissociation and reassociation, leading to specific binding between DNA and protein. In the case of the USS this specific binding then enables uptake of the DNA fragment across the cell’s outer membrane.

The model did not explicitly simulate the first step, non-specific binding, since this is expected to be equally probable for all DNA positions. The specific binding and DNA uptake steps were separately modeled since they are expected to depend on the properties of the DNA uptake machinery and on the length and sequence of the DNA fragment. Although in real cells both steps may depend on the quality of the USS, for simplicity the initial version of the model assumed that specific binding required only a threshold similarity to the USS consensus and that the subsequent probability of uptake depended on the strength of this similarity.

Simulating these steps required first specifying the genomic sequences that should be treated as USS. This was not straightforward because genomes contain many USS variants that differ in how well they promote DNA uptake (Findlay and Redfield, 2009; Maughan and Redfield, 2009). Our strategy was to score the information content (in bits) of every genome position, using the Position-Specific Scoring Matrix (PSSM) from Mell *et al*.’s degenerate-sequence uptake experiment (Mell et al., 2012) (Table S1), and to use overrepresentation of high-scoring sequences as the USS criterion. We scored every genome position in the three *H. influenzae* strains used for the experiments described below (Table 1), and in four randomly generated sequences with the same length and base composition (Figure S1 shows the score distributions). In the *H. influenzae* genomes, overrepresentation of high-scoring sequences was detectable above a score of 7.0 bits and became dramatic above 10.0 bits, where the numbers of high-scoring positions increased in *H. influenzae* genomes but became vanishingly small in the random-sequence controls, (see inset in Figure S1). Since the slight overrepresentation of scores between 7 and 10 bits was hypothesized to be not a direct effect of DNA uptake but an indirect consequence of mutational degeneration of high scoring USSs, the model initially used a USS cut-off score of 10 bits (‘USS10’, n=1941 in strain 86-028NP). This was later reduced to a less stringent 9.5 bits after weak uptake effects had been examined (‘USS_9.5_’, n=2248 in strain 86-028NP).

**Table 1.** Bacterial strains used in this study

| Strain number | Strain name | Phenotype | Source |
| --- | --- | --- | --- |
| RR3117 | *rec2::spec* | Spectinomycin resistant Rd derivative. No translocation of taken-up fragments to the cytoplasm | Sinha *et al.* 2012 |
| RR3125 | *Rd Δrec2* | Unmarked Rd derivative. No translocation of taken-up fragments to the cytoplasm | Sinha *et al.* 2012 |
| RR3133 | 86-028NP<br>NalR | Otitis media clinical isolate. Nalidixic acid resistant | Mell *et al.* 2011 |
| RR1361 | *PittGG* | Nontypeable clinical isolate | G. Ehrlich |
| RR722 | *Rd* | Rd KW20, rough (unencapsulated) derivative of type d | H. O. Smith |

The computational model used these USS scores to predict DNA uptake for every position in the genome, summing the contributions of binding and uptake probabilities from DNA fragments of different sizes (Figure 2A gives an overview). Each fragment under consideration was first checked for the locations of any USS_10_s, and the maximum fragment length (20 kb) and the mean length of USS-free segments (*mean_gap*) were used to calculate the probability that a DNA receptor protein initially encountering a random location in the fragment would then encounter and bind specifically to a USS_10_ rather than disassociating from the fragment: *p_bind = 1 – mean_gap/20000*). Fragments with no USS were initially assigned a baseline *p_bind* of 0.1.

**Figure 2.**
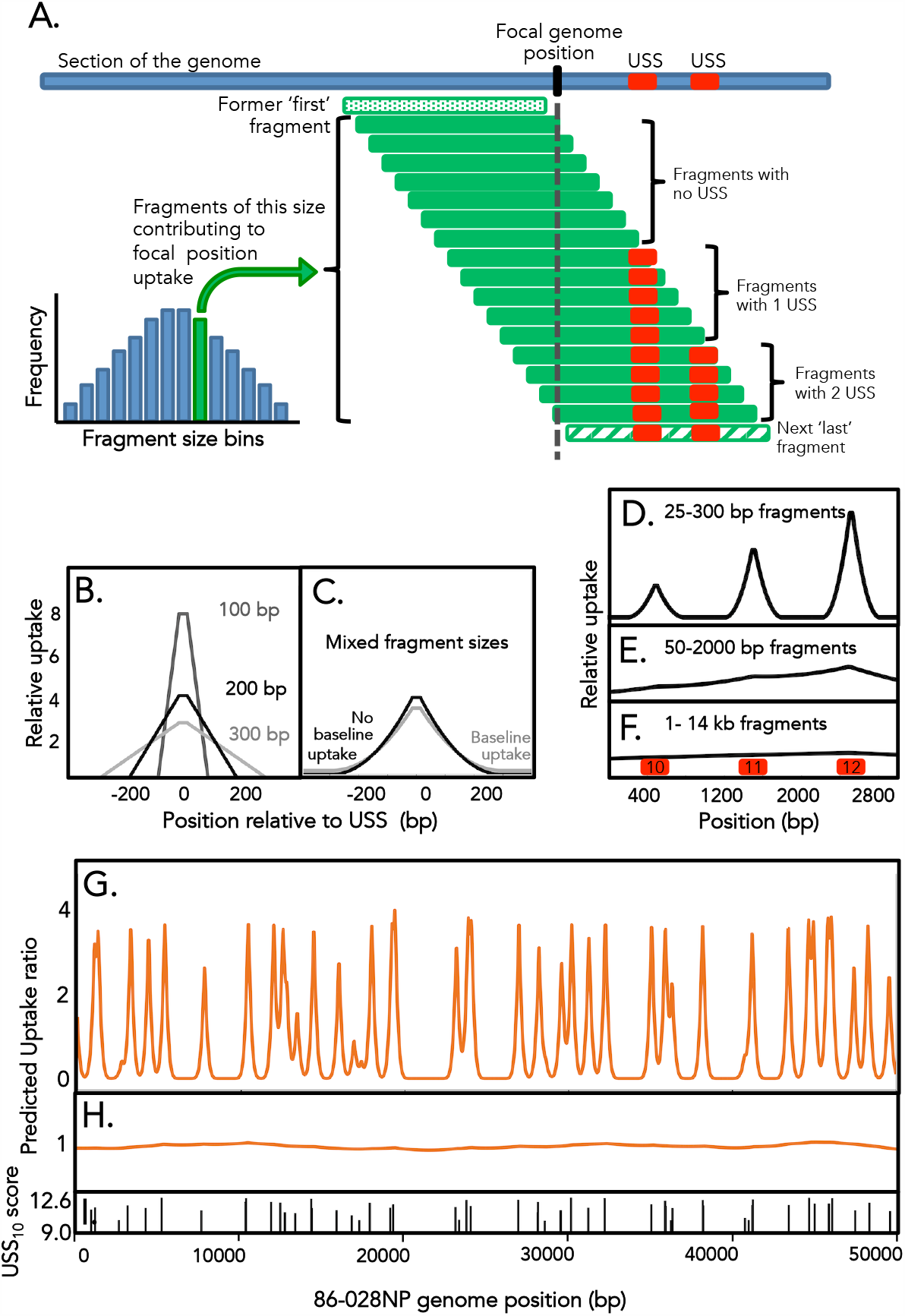
A computational model to predict DNA uptake. **Legend: A**. Components of the DNA uptake model (see Transparent Methods and Results for details). **B. & C**. Model predictions for uptake centered at a 12-bit USS for: **B**. 100, 200, and 300 bp fragments, **C**. a mixed distribution of fragments between 25-300 bp with and without baseline uptake. **D. E. & F**. Model predictions for uptake of a 3000 bp region with 3 USSs (red squares, scores in black) using different fragment-length distributions: **D**. 50-300 bp fragments, **E**. 50-2000 bp fragment, **F**. 1-14 kb fragments. **G. & H**. Predicted DNA uptake of a 50 kb segment of the 86-028NP genome using **G**. short and **H**. long fragment-length distributions: **I**. Locations and scores of USS_10_s in this 50 kb segment. See also Table S1 and Figures S1 and S2.

The probability that this specific binding led to uptake of the DNA fragment was then calculated from the USS_10_ score (or the mean score if the fragment contained more than one USS_10_), using the function *p_uptake = 0*.*1 +* (*1 - 0*.*1*)*/*(*1 + exp*(*-5 \** (*score – 11*))), where *0*.*1* specifies the baseline uptake of USS-free fragments and *-5* is an arbitrarily chosen coefficient specifying the slope at the inflection point. The score of 11 bits specifying the inflection point of the function was chosen because this score was the midpoint of the range of increasing USS overabundance in the genomes (see Figure S1). When combined with the baseline *p_bind* of 0.1, the *p_uptake* baseline assigned fragments with no USS_10_ a net uptake probability of 0.01. Once the contributions of every size of fragment had been calculated for each position (Figure 2A), the model combined all the contributions, taking into account the frequency of each size in the input DNA. The position-specific uptake predictions were then normalized to a genome-wide mean uptake probability of 1.0. Except in simple test cases, for computational efficiency DNA fragment lengths were specified as the median lengths of bins (10 bp bins for short fragments, 200 bp bins for long fragments) rather than each being considered separately (e.g. 101-120 bp, 121-140 bp).

### Model results

Figures 2B and 2C show examples of model predictions for simple situations. Figure 2B shows the uptake predictions for an 800-bp simulated genome containing a single USS with score 12.0 bits, considering three different input DNA fragment sizes (100, 200 and 300 bp). The peaks at the USS have straight sides, a basal width twice the length of the fragments being taken up, and 31-bp flat tops arising from the model’s requirement for a full-length USS. When the DNA fragment sizes were evenly distributed between 25-300 bp in length (Figure 2C), the peak had steep sides at its tops and gradually flattened at the base; maximum width at the base equalled twice the maximum fragment length. In simulated mini-genomes with more than one USS (Figures 2D, E and F), isolated peaks were only seen when the DNA fragments being taken up were substantially shorter than the spacing of the USSs (Figure 2D), and peaks disappeared entirely when the fragments were long enough that almost all contained at least one USS (Figure 2F).

Figure 2G and H show the predicted uptake maps when this model analyzed a 50kb segment of the *H. influenzae* 86-028NP genome, using the short-fragment and long-fragment length distributions from the actual uptake experiments described below (Figures S2A and B), and Figure 2.I shows the distribution of USSs over this segment. Because the ‘short’ DNA fragments are shorter than the typical separation between USSs, the model predicts that uptake will be restricted to sharp peaks at each USS. In contrast, uptake of long DNA fragments is predicted to be much more uniform, since most of these will contain at least one USS.

### Generation of experimental DNA uptake data

To obtain high-resolution measurements of actual DNA uptake, we sequenced *H. influenzae* genomic DNA that had been taken up by and recovered from competent *H. influenzae* cells. Competent cells of the standard laboratory strain Rd were first incubated with genomic DNA preparations from strains 86-028NP and PittGG, whose core genomes are readily differentiated from Rd (and each other) because they differ at ~3% of orthologous positions (Hogg et al., 2007). To allow efficient recovery of the taken-up DNA, the Rd strain in which competence was induced carried a *rec2* mutation that blocks translocation of taken-up DNA fragments, causing the DNA to be trapped intact in the periplasm (16). The 86-028NP and PittGG genomic DNAs were pre-sheared to give short (50-800 bp) and long (1.5-17 kb) DNA preparations (size distributions are shown in Figure S2), and three replicate uptake experiments were done with each DNA preparation. After 20 min incubation with competent cells, the taken-up DNA was recovered from the cell periplasm using the cell-fractionation procedure of Kahn *et al*. (Barouki and Smith, 1985; Kahn et al., 1983; Mell et al., 2012). Taken-up DNA samples were sequenced along with samples of the input 86-028NP and PittGG DNAs and of the recipient Rd DNA. The input and uptake reads were then aligned to the corresponding 86-028NP and PittGG reference sequences and coverage at every position was calculated. Table S2 provides detailed information about the four input samples, the twelve uptake samples, and the Rd sample.

### Removal of contaminating Rd DNA

Preparations of DNA recovered from the periplasm after uptake always included some contaminating DNA from the recipient Rd chromosome. The divergence between the Rd and donor genomes allowed us to estimate the extent of this contamination by competitively aligning the taken-up reads from each sample to an artificial reference ‘genome’ consisting of both recipient and donor genomes as separate ‘chromosomes’. Reads that uniquely aligned to only one chromosome could then be unambiguously assigned to either donor (taken-up) or recipient (contamination). The resulting estimates of Rd chromosomal contamination were between 3.2% and 19.3% of reads; sample-specific values are listed in Table S2.

The effects of this contamination were not expected to be uniform across each donor genome, since segments of the 86-028NP and PittGG genomes with high divergence from or with no close homolog in Rd would be free of contamination-derived reads. We used the competitive-alignment described above to create contamination-corrected uptake coverages by discarding all reads that could not be uniquely mapped to the donor genome; in addition to removing Rd contamination this also removed reads from segments that are identical between the donor and recipient strains (‘double-mapping reads’) and reads that mapped to repeats, such as the six copies of the rRNA genes. For consistency, the same changes were applied to the input samples although they did not experience any contamination. This correction removed an average of 18.6% of reads (range 8.9%-28.3%), left some segments of the 86-028NP and PittGG genomes with no coverage in all samples (2.3% and 2.1% respectively), and reduced coverage adjacent to these segments. Figure S3 shows the locations of the missing data. Contamination details for each sample are provided in Table S2, and the impacts are considered below. The uptake analysis described below showed this correction to be effective.

### Uptake ratios

To control for position-specific differences in sequencing efficiency, contamination-corrected read coverage at each position in each uptake sample was divided by read coverage at that position in the corresponding input sample (*e*.*g*. coverages of each 86-028NP-short uptake sample were divided by 86-028NP-short input coverages). Normalizing the mean of the three replicates to a genome-wide mean ratio of 1.0 then gave a mean ‘uptake ratio’ measurement for each genome position for each DNA type. Finally each position’s uptake ratio was smoothed using a USS-length (31 bp) window. Figures 2.5 and 2.6 show the resulting uptake ratio maps.

**Figure 5.**
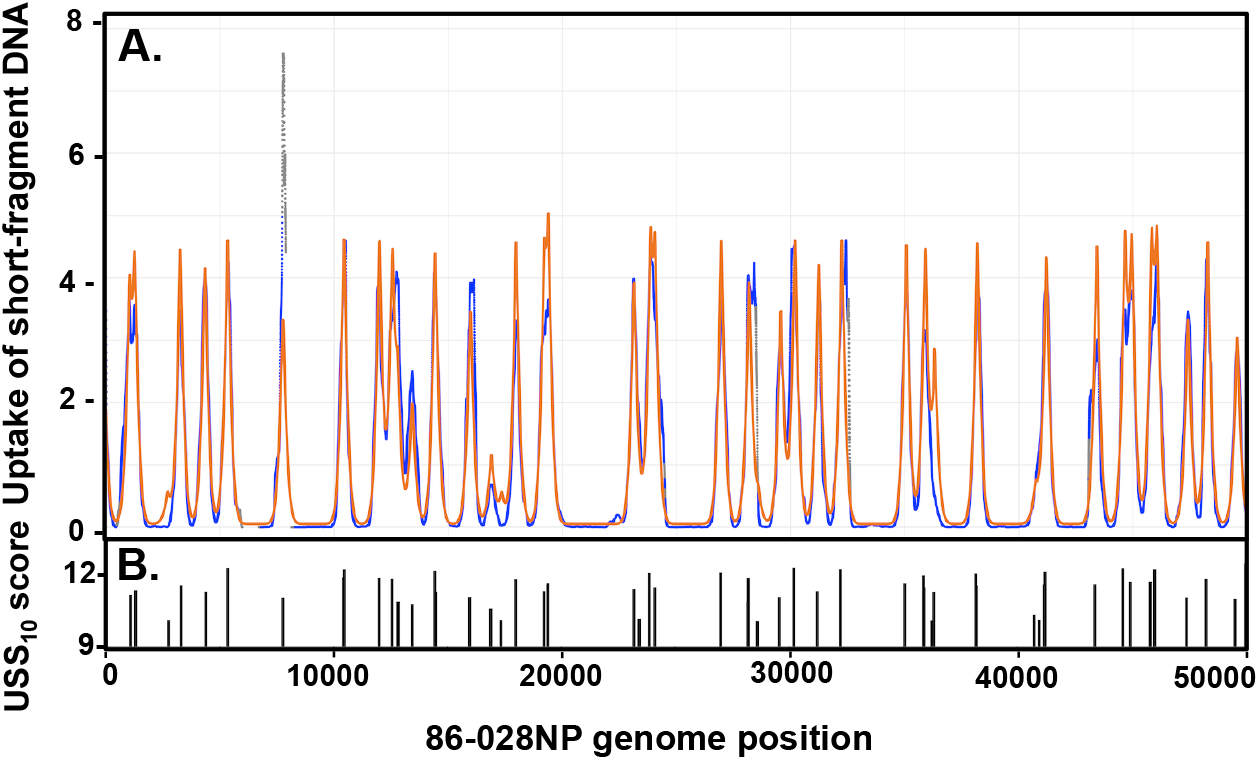
Predicted and observed DNA uptake analysis for short fragments of 86-028NP DNA. **Legend: A**. Map of uptake ratios and initial model predictions. The blue points show the same uptake ratio map as in Figure 3A. The orange points show the same predicted uptake as in Figure 2G. **B**. Locations and scores of USS_9.5_s.

**Figure 6.**
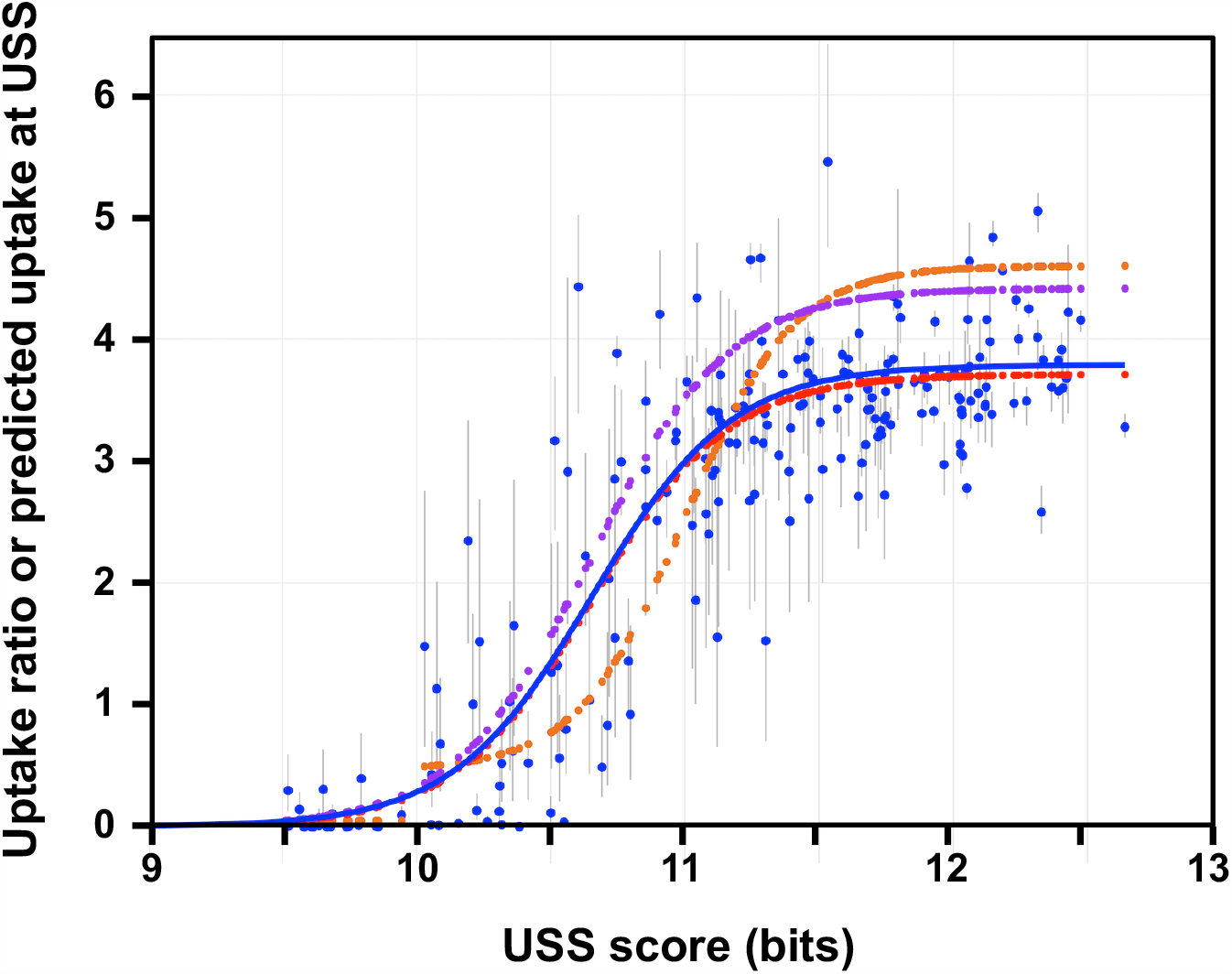
Short-fragment uptake ratios and predicted DNA uptake at isolated 86-028NP USS as a function of USS score. **Legend:** Predicted or measured DNA uptake at 209 USS_9.5_ positions separated by at least 1000 bp from the nearest USS_10_s. The blue dots show the measured uptake ratios; grey bars show the ranges of the three replicates at each position. The blue line shows a sigmoidal function fit to these points. The small orange dots and line show uptake predicted by the original model at the same positions, and the small purple and red dots show uptake predicted by the intermediate and revised model versions discussed in the text. See also Figure S6.

Figure 3A shows the short-fragment uptake ratio map for the first 50kb of the 86-028NP genome; the ticks in Figure 3C indicate locations and scores of USS_10_s. The pattern is strikingly similar to that predicted for the same DNA segment by the model (Figure 2G). Sharp uptake peaks are seen at USS_10_ positions, some separated by flat-bottomed valleys and others overlapping. Figure S4A, B and C show similar analysis for the first 50 kb of strain PittGG’s genome, and Figure S5A and B show expanded maps for two examples of the 86-028NP peaks. The full-genome maps of these uptake ratios are provided in Figure S4D (86-028NP) and S4G (PittGG;) they display the consistency of the peak heights across each genome.

**Figure 3.**
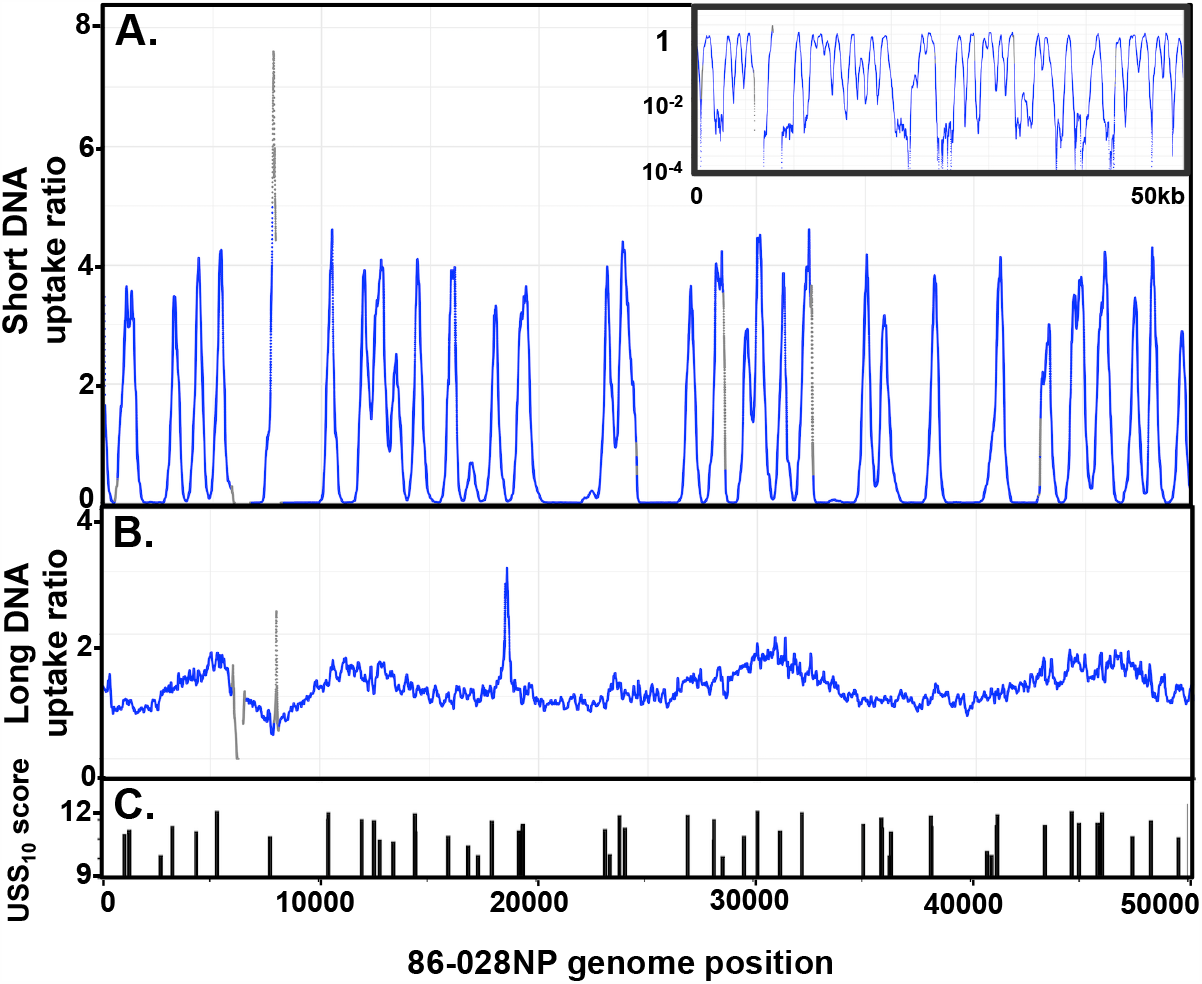
Experimentally determined uptake ratios for a 50 kb segment. **Legend:** The X-axis is the same 50 kb segment of the 86-028NP genome as Figure 2G and H. Grey points indicate positions with input coverage lower than 20 reads. Gaps indicate unmappable segments. **A**. Uptake ratios of short-fragment DNA. **Inset:** Same data with a logarithmic-scale Y-axis. **B**. Uptake ratios of long-fragment DNA. **C**. Locations and scores of USS_10_s. See also Figures S3 and S4.

As expected, the long-fragment DNA samples (Figure 3B and Figure S4B, E and H) had much less variation in uptake ratio than the short-fragment samples; 90% of positions had uptake ratios within two-fold of the mean, and there were few high peaks or low valleys. The few extended segments with low or no uptake coincided with large gaps between USS_10_s. The largest gap was in the 86-028NP segment between 95 and 145kb —the site of a genomic island that is absent from the PittGG and Rd strains, has few USS and has high similarity to an *H. influenzae* plasmid (Harrison et al., 2005). However, uptake ratios did exhibit substantial short-range variation not predicted by the model, which is considered further below.

### Sensitivity is limited by low sequencing coverage

At some genome positions our ability to detect USS-dependent uptake biases and possible USS-independent biases in uptake coverage was limited by low sequencing coverage. Although some of this low coverage arose from the contamination-correction step described above, segments of low coverage were also seen in the raw data, likely arising from biases in the library preparation and sequencing steps. Figure S6 compares the raw coverage of short and long input samples for a 50 kb segment of the 86-028NP genome, illustrating the strong variation in sequencing coverage that was both broadly reproducible and sequence dependent. Low coverage had similar effects in all samples, precluding calculation of uptake ratios where input coverage was zero and generating high levels of stochastic variation where coverage was low (segments with no coverage and positions with coverage below 20 reads are indicated in uptake ratio maps by gaps and grey dots respectively).

We used the low uptake ratios of positions at least 1000 bp from the nearest USS_9.5_ to assess the effectiveness of the contamination correction described above, since these positions are where contamination would have had its largest effects. This peak-separation distance was chosen because a peak-shape analysis of uptake around 158 USS_10_s that were separated by at least 1200 bp from other USS_10_s and had uptake ratios of at least 3.0 found that mean uptake ratio had fallen to baseline at positions 600 bp from the USS (Figure S7). Figure 4 compares uptake ratios between the valley positions that received the contamination correction described above (those with a Rd homolog) and the positions where no correction was needed (those with no Rd homolog). The two distributions had nearly identical medians (0.0022 and 0.0020 respectively), indicating that the correction was sufficient but not excessive. These low values also confirm that the DNase I treatment and washing steps removed at least 99.99% of the donor DNA that had not been taken up.

**Figure 4.**
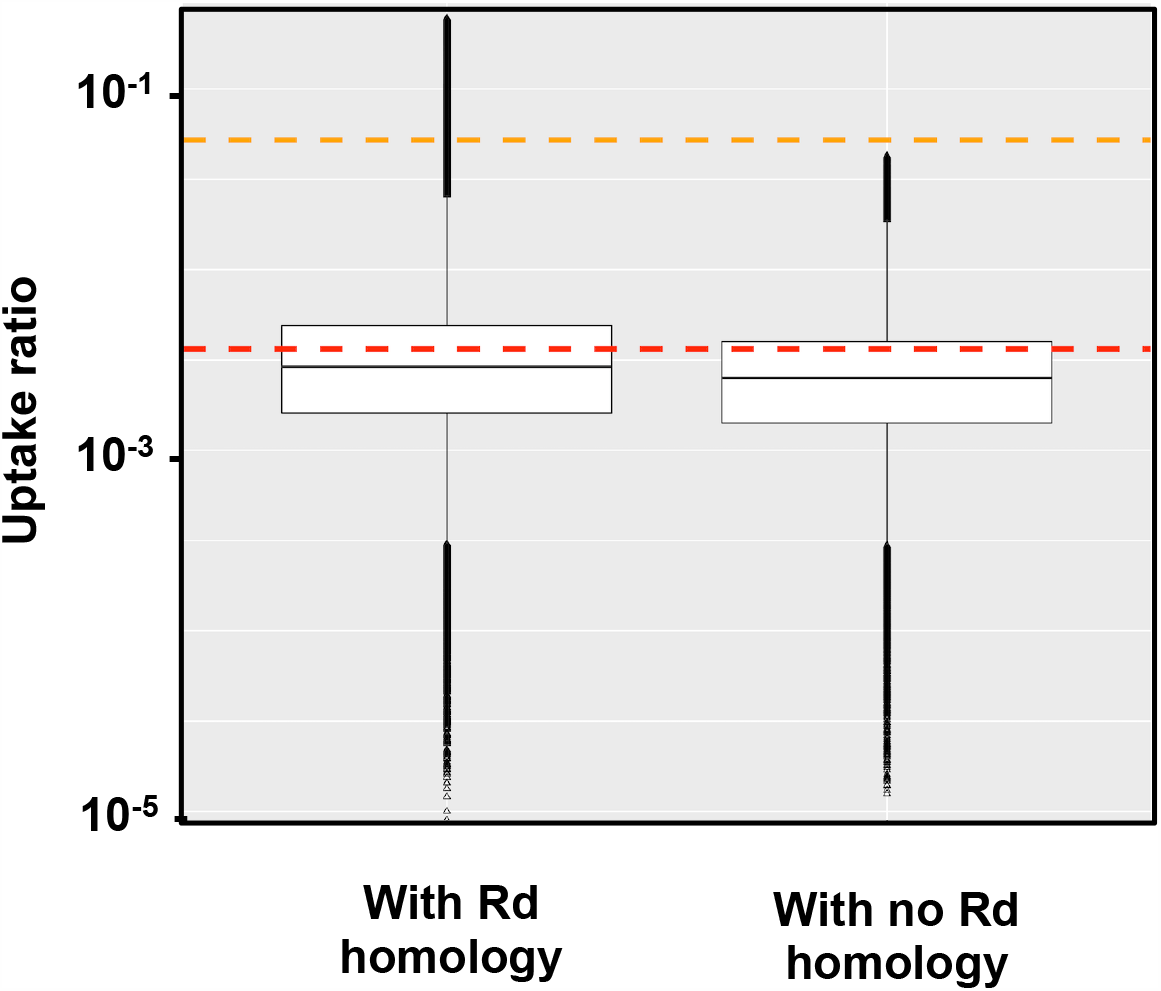
Depths of uptake ratio valleys with and without contamination correction. **Legend:** Uptake ratios of positions in the 86-028NP short-fragment dataset that were at least 1kb from the nearest USS_9.5_ and whose uptake coverages were either corrected for contamination with homologous Rd sequences (left, 272,547 positions) or not corrected for contamination because they had no Rd homology (right, 89,418 positions). Orange and red dashed lines indicate valley uptake predicted by the original and revised models respectively. See also Figure S6.

### Uptake ratios show no periodicity across the genome

Bacterial genomes show periodicity for several features related to DNA curvature and codon usage biases (Mrazek, 2010), so we examined the distribution of uptake ratios across each genome by Fourier analysis, using the R package TSA. The log-log plots in Figure S8 show that this found no strong influence of any specific repeat period on either the variation in input-sample coverage (panels A-D) or the variation in uptake ratios (panels E-H). Instead, to explain the observed variation, the analysis needed to invoke small contributions from almost every possible repeat period.

### Uptake of short-fragment 86-028NP DNA

To investigate how USSs contribute to the DNA uptake process, our strategy was to first analyze discrepancies between model predictions and observed uptake ratio peaks in the 86-028NP short-fragment dataset, since these would reveal ways in which the simple assumptions underlying the model mischaracterized the actual steps of DNA uptake. Model changes that improved the predictions were considered to better reflect the true constraints on uptake of short DNA fragments. The refined model’s predictions were then compared to the real uptake ratios for 86-028NP long-fragment DNA, allowing additional refinement, and finally to the short- and long-fragment uptake ratios for PittGG DNA.

Figure 5 compares predicted and measured uptake (orange and blue lines respectively) of short-fragment 86-028NP DNA for the first 50kb of the genome. The close correspondence between the model’s predictions and observed peak locations and shapes confirmed that almost all of the variation in uptake was due to USSs. (The Pearson correlation coefficient over all positions was 0.92). However, the more detailed analyses below identified components of the model that could be improved.

First, the predicted low-uptake valleys were too high. This is not easily seen in Figure 5, but the log-scale inset in Figure 3 and the analysis in Figure 4 show that most valley-bottom positions had uptake ratios between 0.01 and 0.001, well below the model’s predicted baseline uptake of 0.052 (dashed orange line in Figure 4).

Second, the score cutoff of 10.0 bits was too high. Analysis of score-subsets within the uptake-valley dataset showed that 9.5 bits was a better cutoff. For the 78 positions with scores between 9.5 and 10.0 bits, the correlation between score and uptake ratio was 0.27. Although a similar correlation (0.27) was seen for the 136 positions with scores between 9.0 and 9.5 bits, it was driven mainly by very small effects, and only 4 positions had uptake ratios higher than 0.1. For the 461 positions with USS scores between 8.5 and 9.5 bits the correlation was only 0.09.

Third, most of the predicted peaks were too high. Figure 6 complements Figure 4’s analysis of uptake valleys with an analysis of similarly isolated uptake peaks. The blue dots show the uptake ratio at every USS_9.5_ position in the genome that is at least 1000 bp from the nearest USS_10_ (n=209), and the small orange dots show predicted uptake at the same positions. The lack of scatter in the orange points confirms that the separation distance was sufficient to avoid predicted effects of nearby USS; the least-squares difference between predicted and observed uptake ratios at these 209 USS positions was 0.521.

In the original-version of the uptake model, the sigmoidal parameters of the *p_uptake* function (baseline and location and slope of inflection point) were set using the frequencies of USS scores in the 86-028NP genome; as expected, the same parameters were obtained when a sigmoidal function was fit to the uptake points predicted by this model (orange line in Figure 6). To improve the predictions we replaced these *p_uptake* parameters with those of a sigmoidal function fitted to the real uptake data (blue line in Figure 6), changed the *p_uptake* baseline to 0.005, lowered the USS score cutoff from 10.0 bits to 9.5 bits, and ran the model again. The purple points in Figure 6 show that, although peak-height predictions for low-scoring USS were somewhat improved by the changed *p_uptake* function, those for USS with scores > 11.5 bits became slightly worse; the least-squares difference for the 209 positions was modestly reduced from 0.521 to 0.420.

We initially suspected that the over-prediction of peak heights might be due to overestimating the proportion of very short fragments in the input DNA, but eliminating the contributions of fragments shorter than 80 bp had very little effect on predicted peak heights. (This is likely because these fragments contribute little DNA and rarely contain USS.) We then considered an alternative explanation, that USS might be ineffective when they were very close to fragment ends. The red dots in Figure 6 shows that requiring USS to be at least 50 bp from fragment ends lowered predicted peak heights to the mean observed height while maintaining a low baseline uptake. Distances of 30 bp and 70 bp were also tested, but gave peaks that were, respectively, too high and not high enough. Incorporating this modification into the model barely changed the least-squares difference of its predictions with the observed uptake ratios (0.93, up from 0.92), but dramatically reduced the least-squares difference from 0.420 to 0.052. This revised model was used by the analyses described below.

The above analyses do not explain why many uptake ratio peaks were substantially higher or lower than predicted by their USS score. The scatter is unlikely to be due to noise alone, since more extreme smoothing of the uptake ratio data with windows as large as 150 bp improved the correlation by only 0.001. Below we consider three other factors that might influence DNA uptake: weak non-USS uptake biases, effects of interactions between bases at different positions in the USS, and effects of DNA shape.

Weak uptake biases could arise from either low-scoring USS or non-USS sequence factors. Since weak biases would only be detectable in genome segments that lack strong USSs, we searched for them using a far-from-USS_10_ dataset containing only DNA segments whose ends were at least 0.6kb from the closest USS_10_ and that had input coverage of at least 20 reads. This dataset contained 513 segments where weak uptake effects could in principle be detected (22% of the genome); their mean uptake ratio over the 428678 positions was 0.0102. In these segments, the only positions with distinct uptake ratio peaks >0.2 were 9 weak USSs with scores between 9.5 and 9.9 bits (an example is shown in panel C of Figure S5). Since this analysis did not find any non-USS positions giving uptake higher than 0.2, it suggests that other sequence factors do not detectably promote uptake in the absence of a USS.

The degenerate-USS analysis of Mell *et al*. (Mell et al., 2012) found that pairwise interactions between the AT-tract bases and core bases of a USS made substantial contributions to uptake of a 200 bp synthetic fragment. To evaluate the effects of these interactions in predicting uptake of genomic DNA, the USS scores used by our revised model were raised or lowered in proportion to the interaction effects reported in Figure 6 of Mell *et al*. Figure 7 shows that, for the same 209 isolated USS_9.5_s used above (Figure 6), this adjustment had little effect on high-scoring USS but further reduced the scores of low-scoring USS. The data points are coloured by their uptake ratios, showing that weak USS giving unusually high or low uptake were equally likely to have their scores reduced. However, these scoring changes had no effect on the short-fragment uptake predictions of the revised model (both Pearson correlations 0.93), probably because (i) the scores of strong USS were not significantly changed and (ii) the scores of weak USS already gave near-baseline uptake.

**Figure 7.**
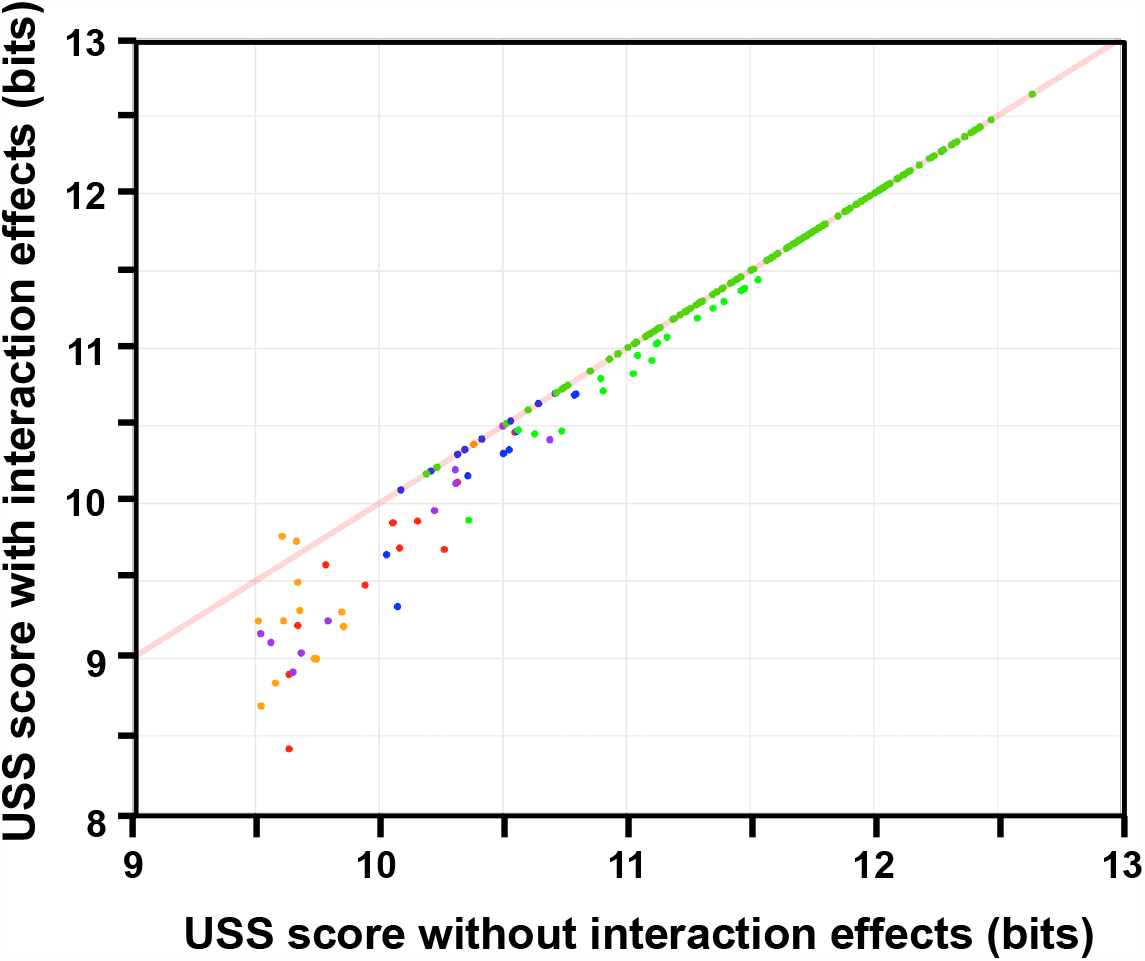
Effects of within-USS interactions on USS scores. **Legend:** USS scores calculated with and without interactions effects for 209 isolated 86-028Np USS_9.5_s (see Methods for details). Red line shows expected scores if the interactions had no effect. Point color indicates the uptake ratios at the USS: yellow, <0.01; red, 0.01-0.1; purple, 0.1-0.5; blue 0.5-1.5; green, >1.5.

### DNA shape effects

Most of the interactions in Figure 7 were between bases separated by 10 bp or more, so we complemented that analysis with analysis of DNA shape, which reflects both pairwise and more complex interactions over a 5 bp range. The major shape features that can be predicted from DNA sequence are the minor groove width, the propeller twist between two paired bases, the helix twist between one base pair and the next, and the roll of one base pair relative to the next. In Figure 8, the wide grey line reproduced in each panel shows these feature predictions for the consensus USS. The USS inner core (orange shading) has a relatively wide minor groove and high propeller twist, which would facilitate sequence recognition by proteins (Rohs et al., 2009). To the left of this and in both AT-tracts (yellow shading) the minor groove is narrow with low propeller twist and negative roll, predicting that these segments are both rigid and slightly bent.

**Figure 8.**
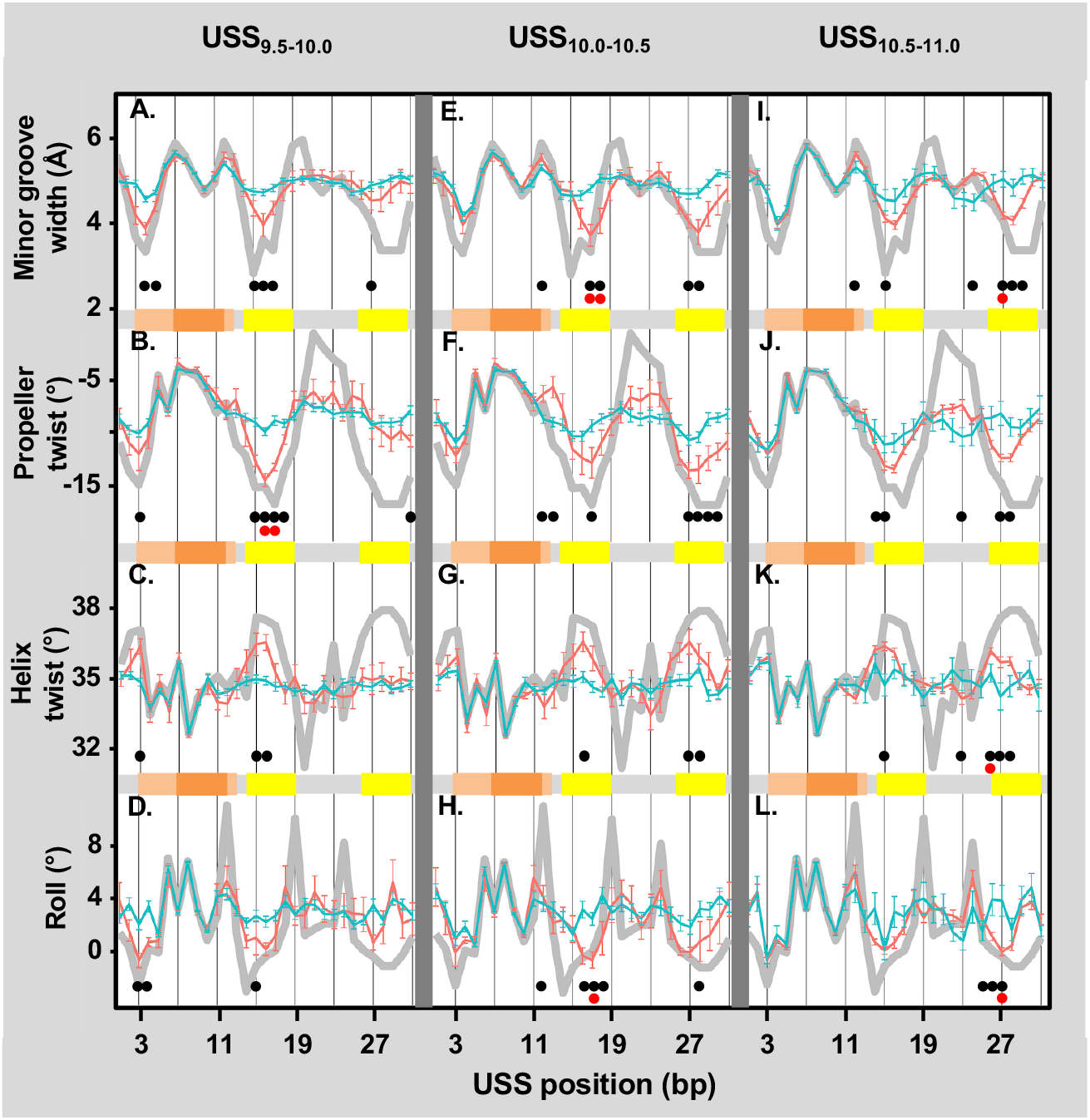
Predicted shape features of USS. **Legend:** The thick grey line reproduced in each box shows shape analysis of the consensus USS sequence. Blue and orange lines in each box show shape analysis of genomic USS separated by at least 500 bp, grouped by score and coloured by uptake ratio. The orange and yellow bars below each box indicate components of the USS (see Figure 1): light orange: outer core; dark orange: inner core; yellow: AT tracts. **Panels A-D** USS_9.5-10_: Blue: uptake ratios <0.2 (n=68, mean USS score =9.7). Orange: uptake ratios >0.2 (n = 10, mean USS score =9.8). **Panels E-H:** USS_10.0-10.5_: Blue: uptake ratios <0.6 (n=47, mean USS score=10.22). Orange: uptake ratios >2.0 (n=10, mean USS score=10.26). **Panels I-L:** USS_10.5-11.0_: Blue: uptake ratios <0.6 (n=14, mean USS score=10.64). Orange: uptake ratios >2.0 (n=59, mean USS score=10.79). **A, E and I**. Minor groove width, in Å. **B, F and J**. Propeller twist, in degrees. **C, G and K**. Helix twist, in degrees. **D, G and L**. Base pair roll, in degrees. See also Figure S6. Grey bars for each point show the standard error. Dots indicate significant differences by Kolmogorov-Smirnov between high-uptake and low-uptake positions with (red dots) and without (black dots) Bonferroni correction.

To see if shape features affect uptake in ways that were not captured by scores alone, we compared the shape features of subsets of the 209 isolated USS with similar scores but different uptake ratios. Panels A-D of Figure 8 compare the shape features of very weak USS (USS_9.5-10_) whose uptake ratios were either low (<0.6, blue lines) or high (>2.0, orange lines). Similarly, panels E-H and I-L show the same comparisons for USSs with low and moderate scores (USS_10-10.5_ and USS_10.5-11_ respectively). USSs with scores higher than 11 bits were not analyzed since they did not exhibit enough uptake variation to reveal correlations between uptake and DNA shape.

In all three score subsets the inner-core shape features were very similar for low-uptake and high-uptake subsets (blue and orange lines), probably because this sequence perfectly matches the consensus in 206 of the 209 USS_9.5_. (The other three had otherwise-perfect core sequences and good AT tracts but had only baseline uptake ratios.) However, the AT-tract shapes had marked differences, with low-uptake USSs having no distinctive shape features and high-uptake USSs resembling the USS consensus shape. This suggests that the predicted rigidity and slight bend caused by interactions within the consensus AT tracts facilitate DNA uptake.

### Uptake of fragments with more than one USS

Many genomic USSs are sufficiently close that they will co-occur even on short DNA fragments; 23% of 86-028NP USS_10_s are within 100 bp of another USS_10_, and 17% are within 30 bp (Figure S9A). Fragments with multiple USS might be expected to have relatively high uptake, since they provide more targets to which the uptake machinery receptor could bind, but only one of the two previous studies in *Neisseria* found this effect (Ambur et al., 2007; Goodman and Scocca, 1991).

Visual examination of uptake ratio maps around the 230 pairs of 86-028NP USS_10_s within 100 bp found only single peaks; Figure S9B shows that the midpoints of these USS pairs (red and green points) have very similar uptake ratios to those of the 209 isolated USSs (pale blue points). Figure S5 panels E and F show examples of peaks at USS pairs separated by 69 bp and 230 bp.

A special subclass of USS pairs consists of oppositely oriented pairs that are so close that they overlap; these can form RNA hairpins and are usually located at the ends of genes where they act as transcriptional terminators (Kingsford et al., 2007; Smith et al., 1995, 1999). Figure S9A shows that the 86-028NP genome has 109 USS_10_ pairs whose centers are within 14 bp: 69 in the −/+ orientation, all 0-3 bp apart, and 40 in the +/- orientation, all 10-14 bp apart. The green points in Figure S9B shows that midpoint uptake ratios of these pairs were very similar to those of other pairs or at isolated USS_10_s with similar scores.

Overall, these results indicate that the presence of two USS_10_s within 100 bp does not detectably increase the probability of the receptor binding to a USS, a result consistent with that of Ambur *et al*. for pairs of closely spaced DUS in *Neisseria meningitidis* (Ambur et al., 2007). Because individual fragments were not tracked, we cannot make any conclusions about uptake of fragments with more widely separated USS.

### Uptake of long-fragment 86-028NP DNA

Figure 9A compares the revised model’s predictions for long-fragment 86-028NP DNA with the uptake ratios observed over the same 50kb genome segment as in Figure 3B. In contrast to predictions for short-fragment uptake (correlation of 0.93), it seriously underpredicted the variation in long-fragment uptake ratios (correlation of 0.60). Although 51% of 86-028NP positions had long-fragment uptake ratios lower than 0.8 or higher than 1.2, only 13% had predicted uptake outside these limits.

**Figure 9.**
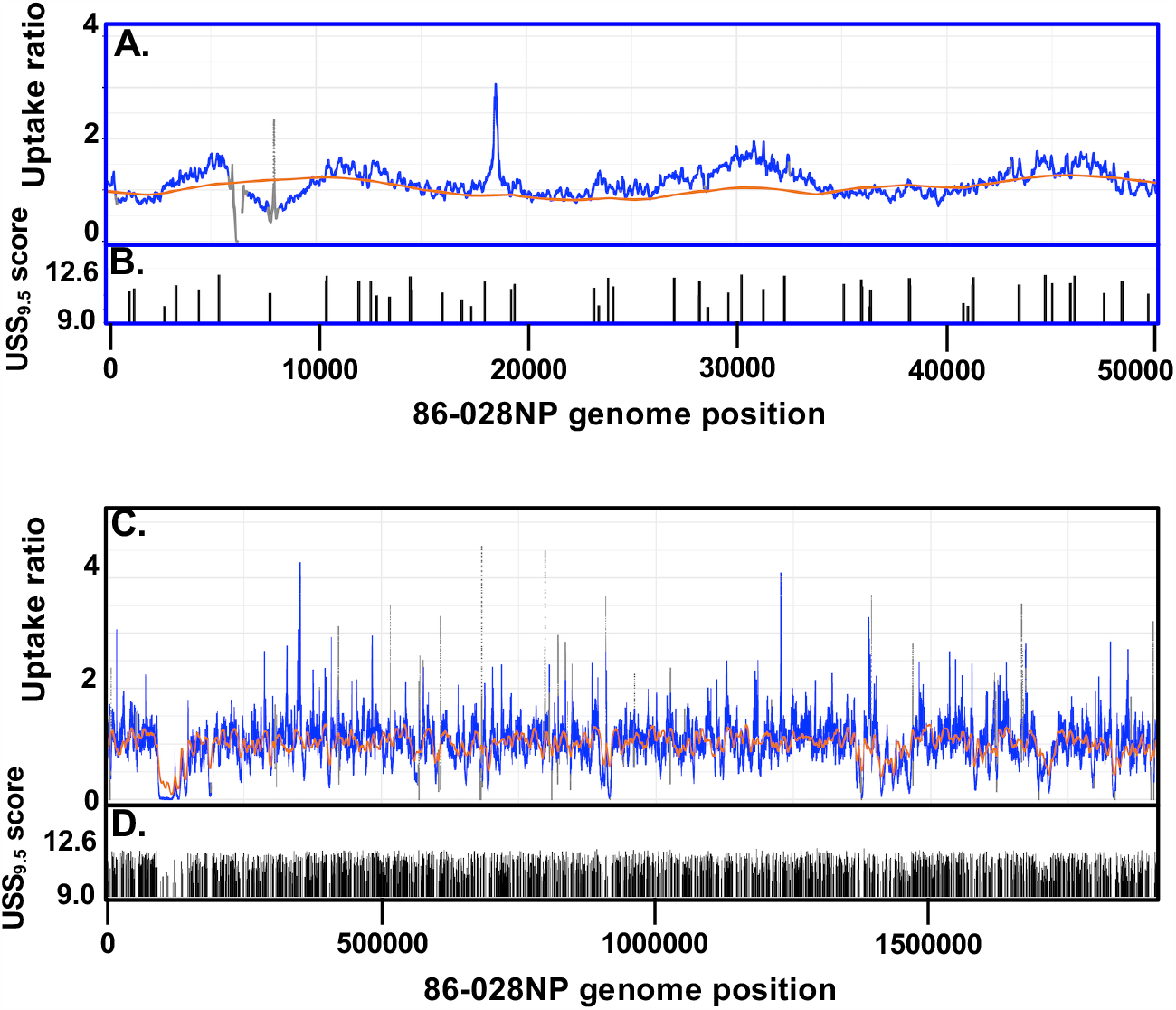
Predicted and observed uptake of long 86-028NP DNA fragments. **Legend: A. and C**. Uptake maps for 86-028NP long-fragment DNA. Orange points: USS-dependent uptake predicted by the revised model. Blue points: mean uptake ratios from 3 replicate experiments (grey indicates input coverage <20 reads, gaps indicate unmappable positions). **B**. and **D**. USS_9.5_ positions and scores. **A. and B**. The same 50 kb genome segment shown in previous figures. **C. and D**. Whole genome.

Because long-fragment uptake ratios lacked the dramatic peaks and valleys seen for short-fragments, stochastic noise arising at the regions of low sequencing coverage described earlier was expected to play a larger role. To estimate the magnitude of this effect, we compared the effects of adding different amounts of artificially generated noise to simulated (noise-free) uptake data (Figure S10A shows examples of noise-free and noise-added coverages). Figure S10B shows that, as expected, the correlation between noisy and noise-free data worsened as the arbitrary level of noise increased for both short-fragment (blue) and long-fragment (red) simulations, and that increasing noise had a much stronger effect on the long-fragment simulations. For short DNA fragments, simulations with noise levels of 2 and 2.5 gave correlations of 0.94 and 0.92, very close to the 0.93 correlation between the revised model and the real data. (dashed blue line in Fig. S9B) For long DNA fragments with the same noise levels, more noise was needed, with a 3.0 multiplier giving a correlation of 0.56, slightly lower than the observed correlation of 0.60 (dashed red line in Fig. S9B). This confirms that the disparities between measured uptake ratios and USS-based predictions were at least partially due to noise in the sequencing data. However, these correlations are still 21% and 10% higher than the best correlation obtained between the revised model’s predictions and real data.

One notable feature of the long-fragment uptake ratio maps in Figure 3B and 9A is the presence of occasional spikes of unusually high uptake, *e*.*g*. at position 18,540 (examples are shown in panels G and H of Figure S5). These spikes were much narrower than expected for true uptake biases acting on long DNA fragments, so they were likely due to stochastic differences between input and uptake coverage in regions of low coverage, not to true differences in uptake. Consistent with this explanation, 70.4% of positions with uptake ratios greater than 2.0 had coverage less than 100 reads, compared to only 8% of positions with more typical uptake ratios between 0.5 and 2. Table S3 shows analysis of the distribution of uptake extremes at positions with different levels of sequencing coverage, and panels I and J of Figure S5 show the low coverage at the example spikes. However, this explanation did not apply to most positions with very low uptake, which were in broad segments with few or weak USS, not in narrow spikes at low coverage positions, and thus likely reflect genuinely low uptake.

### Prediction of uptake of PittGG DNA

Since the uptake model was refined using uptake data for DNA of strain 86-028NP, the revised model’s predictions were assessed using uptake data for DNA of a different strain, PittGG, which differs from 86-028NP DNA by SNPs and indels affecting about 11% of its genome. Figure S11 compares the uptake predictions with the observed PittGG uptake ratios. For short-fragment data the peak heights and valley depths were similar to those for 86-028NP, as was the Pearson correlation between predicted and observed uptake (0.92). Although the model predicted similar long-fragment variation for 86-028NP and PittGG, the PittGG experimental data showed more extreme variation, and the correlation was only 0.41, substantially worse than the 0.60 obtained for 86-028NP. As with the 86-028NP data, much of this discrepancy may be due to noise in regions of low sequencing coverage. However this may not fully explain PittGG’s lower correlation, because input samples of the two strains had similar frequencies of low-coverage positions (Table 2.2).

**Table 2.** Frequencies of *Hin*-type USS in genomes of other species

| Species | Genome size | USS<sub>10</sub>/Mb | USS<sub>11</sub>/Mb |
| --- | --- | --- | --- |
| *H. influenzae* 86-028NP | 1.914 | 1014 | 754 |
| *H. parainfluenzae* T3T1 | 2.087 | 867 | 683 |
| *Aggregatibacter actino-*<br>*mycetemcomitans* VT1169 | 2.129 | 970 | 737 |
| *H. ducreyi* 35000HP | 1.699 | 102 | 16 |
| *Mannheimia haemolytica*<br>M42584 | 2.732 | 96 | 18 |
| *N. meningitidis* | 2.272 | 52 | 4 |
| *Streptococcus pneumoniae* | 2.039 | 16 | 1 |
| *Pseudomonas aeruginosa* | 6.264 | 8 | 0 |
| *Homo sapiens*<sup>1</sup> | 3 x 1.9 Mb | 6.6 | 0.1 |
| Random-sequence DNA<sup>2</sup>,<br>41% G+C | 3 x 1.9 Mb | 29.5 | 0.3 |

1. Means for three 1.9 Mb segments of human chromosomes 1, 3 and 12.
2. Means for three 1.9 Mb 'genomes' with the same base composition as human
DNA.

### Predicted competition with other DNAs in the human respiratory tract

*H. influenzae*’s natural environment is the human respiratory tract, where its DNA must compete for uptake with DNAs from other bacteria, and with host-derived DNA whose concentration in respiratory mucus of healthy individuals can exceed 300 µg/ml (Lethem et al., 1990; Shak et al., 1990). The revised model was used to investigate this competition.

Table 2 lists the frequencies of USS_10_ and USS_11_ in the genomes of various species. Pasteurellaceae species that share *H. influenzae*’s *Hin*-type USS (*e*.*g. H. parainfluenzae* and *Aggregatibacter actinomycetemcomitans*) typically have about 1000 USS_10_s per Mb (Redfield 2006); most of these are strong USS with scores ≥ 11. Pasteurellaceae with the *Apl*-type USS (*e*.*g. H. ducreyi* and *Mannheimia haemolytica*) have about tenfold fewer *Hin*-type USS_10_s, and fewer than 20% of these are strong. Other common respiratory tract bacteria (*e*.*g. N. meningitidis, Streptococcus pneumonia* and *Pseudomonas aeruginosa*) have USS only at the frequencies expected for their base compositions. The human genome is exceptional in having about 5-fold *fewer* USS_10_ per Mb than expected from simulated sequences with the 41% GC content of human DNA (observed frequency 4.6/Mb, simulated frequency 30/Mb, both with mean scores of only 10.3 bits). Since the USS inner-core motif includes a CpG, the underrepresentation of USS_10_ in human DNA relative to simulated sequences is probably a consequence of the 4-5 fold depletion of CpGs in the human genome caused by deamination of methylated cytosines (Babenko et al., 2017; Bird, 1986).

To estimate how strongly these DNAs would compete with *H. influenzae* DNA for uptake by *H. influenzae* cells, we concatenated the 86-028NP genome with each of two Pasteurellacean genomes and with three 1.9 Mb segments of the human genome, and used the revised uptake model to predict relative uptake. The respiratory pathogen *H. parainfluenzae* was used as the *Hin*-type USS representative. The genital pathogen *H. ducreyi* was used as a representative species with *Apl*-type USSs, since none occur in the human respiratory tract, and segments of human DNA also served to represent non-Pasteurellacean bacteria. To approximate the lengths of DNA fragments in the respiratory tract (Lethem et al., 1990; Shak et al., 1990), the model was run using fixed fragment lengths of 1kb and 10kb, and because human DNA will contain many fragments lacking USSs, the predictions were made with and without the model’s *p_uptake* baseline binding probability of 0.005. For each genome in the concatenated sequence, the predicted uptake at every position was then summed to get a total genome uptake value for each fragment length and baseline assumption.

Table 3 shows the relative amounts of *H. influenzae* DNA predicted to be taken up under the various competition conditions. As expected from published uptake-competition experiments (Albritton et al., 1984; Redfield et al., 2006), *H. influenzae* and *H. parainfluenzae* DNAs were taken up with equal efficiency in all simulated conditions, and *H. ducreyi* DNA was much less competitive, contributing only 5% of the 1 kb fragments and only 12% of the 10 kb fragments. With human DNA as the 1:1 competitor, more than 99% of the DNA taken up was predicted to be from *H. influenzae*. For all competing DNAs, baseline uptake of fragments containing no USS made only a tiny contribution.

**Table 3.** Predicted relative uptake^1^ of *H. influenzae* DNA in simulated competition with DNAs of other species^2^.

| **Assumptions:** | **1 kb**<br>**fragments,**<br>***p_uptake*=0** | **1 kb**<br>**fragments,**<br>***p_uptake***<br>**=0.005** | **10 kb**<br>**fragments,**<br>***p_uptake*=0** | **10 kb**<br>**fragments,**<br>***p_uptake***<br>**=0.005** |
| --- | --- | --- | --- | --- |
| **86-028NP DNA in**<br>**competition with:** |  |  |  |  |
| ***H. parainfluenzae***<br>**DNA (*Hin*-type USS)** | 0.500 | 0.500 | 0.475 | 0.475 |
| ***H. ducreyi* DNA**<br>**(*Apl*-type USS)** | 0.952 | 0.951 | 0.884 | 0.883 |
| ***Homo sapiens* DNA<sup>3</sup>**<br>**(no USS enrichment)** | 0.998 | 0.997 | 0.991 | 0.990 |

<sup>1</sup>. *H. influenzae* DNA as a fraction of the total DNA predicted to be taken up.
<sup>2</sup>. Competitions were genome:genome
<sup>3</sup>. Means for the three genome 1.9 Mb segments described in Transparent Methods.

## DISCUSSION

DNA uptake by competent *H. influenzae* Rd cells was measured at every position in the genomes of two divergent *H. influenzae* strains, using short-fragment and long-fragment DNA preparations. Differences between observed uptake and that predicted by a computational model of USS-dependent uptake revealed the strength of the uptake machinery’s bias towards USS and the absence of other sequence biases. These findings increased our understanding of DNA uptake bias and its potential effects on the distribution of recombination.

### Implications for DNA uptake

#### The USS motif

The measured discrimination’ for USS was very strong; with short DNA fragments, valleys at USS-free segments had ~1000-fold lower uptake ratios than peaks at high-scoring USS. This non-zero baseline is unlikely to be due to residual contamination by recipient DNA, since valley depths were similar for segments with and without Rd homology (Figure 4).

One surprising finding of Mell et al.’s (2012) work was the difference between the inner-core uptake motif identified by their degenerate-USS experiments (Figure 1A) and the extended motif of USS sequences in the *H. influenzae* genome (Figure 1B). Our analysis confirmed that uptake absolutely requires a perfect match to the inner core, but found that this was not sufficient to raise uptake above baseline, even if the rest of the core was perfectly matched.

#### Effect of USS shape

The predicted shape differences between similarly-scoring USSs that gave strong or weak uptake (Figure 8) suggest a preference for USS that are rigidly bent at AT-tracts and the outer core (Harteis and Schneider, 2014; Rohs et al., 2009). Similar preferences have been described for several DNA binding proteins and have been associated with specific binding by arginine or lysine residues to narrow minor grooves (Rohs et al., 2009; Stella et al., 2010). These features have been integrated successfully in some transcription factor binding models (Li et al., 2017), but using them to improve uptake prediction will require more comprehensive investigation into the effects of DNA shape on uptake. The stiffness also suggests that the initial passage of DNA through the secretin pore may be facilitated by transient strand melting at or beside the USS rather than by bending (Danner et al., 1982).

#### Other USS effects on DNA uptake

Comparison of predicted and measured heights of uptake peaks suggested that USSs were ineffective when located very close to fragment ends. This would be consistent with experimental evidence that uptake initiates internally rather than at fragment ends(Barany et al., 1983), but would need to be experimentally investigated. The presence of two USSs within 100 bp did not detectably increase uptake, a finding consistent with Ambur *et al*.’s (Ambur et al., 2007) study of very close uptake sequences in *Neisseri*a.

#### Lack of USS-independent effects

The very low valleys between short-fragment uptake peaks allowed us to examine more than 400,000 positions for USS-independent increases in uptake. None were found; the only positions with distinct uptake ratio peaks >0.2 were 9 weak USSs.

#### Implications for genetic exchange

Preferential uptake of USS can create variation in recombination at all levels: across a single genome, between strains of one species, and between both closely related and unrelated species. Pifer & Smith showed that transformation frequency in *H. influenzae* decreased exponentially when fragments were smaller than 3.5 kb. This decrease was attributed to exonuclease degradation of fragments from their 3’ ends, since similar numbers of short and long fragments were taken up, only 5’-end label was incorporated into the chromosome, short fragments transformed more efficiently when the selected marker was far from one end. Mell (2014) used genome sequencing of inter-strain recombinants to examine the distribution of recombination tract lengths; despite the presence of 2-3% SNPs, the mean tract length was 6.9 kb and the longest was 43 kb. The efficient uptake and recombination of long fragments allows fragments containing non-homologous segments to still recombine well, provided both fragment ends are homologous. Fragments with only one homologous end recombine much less efficiently (‘homology-facilitated recombination’ (de Vries and Wackernagel, 2002)), and integration of fragments with no homology is very rare.

#### Across the genome

Across the genome of a *H. influenzae* strain, the genetic consequences of USS-dependent DNA uptake depend on USS locations and on the lengths of the available DNA fragments. If only short fragments are available, the limitation to positions close to a USS may be obscured by limitation caused by degradation of incoming DNA in the cytoplasm. If most fragments are long, recombination will be both more frequent and more evenly distributed across the genome, because long fragments are more likely to both contain USS and recombine. The result will be that almost all recombination is caused by fragments long enough to usually contain at least one strong USS. This situation is caused by the high abundance and relatively even distribution of genomic USS, itself an expected consequence of the functionally neutral accumulation of USS under a molecular drive caused by USS-biased DNA uptake (Danner et al., 1982; Maughan et al., 2010).

#### Recombination between *H. influenzae* strains

On average about 85% of the genomes of *H. influenzae* strains are homologous, with sequence divergence low enough to have only a modest effect on recombination frequencies (Mell et al., 2011). On average, genome segments that are absent from other strains have lower density of USS (0.58/kb vs 1.08/kb for sequences present in 86-028NP but absent from Rd. Only some of these will be segments newly acquired from species without USS. If a non-homologous segment introduced into one strain by conjugation or transduction is beneficial, the USSs in adjacent DNA will help it efficiently spread to other strains by transformation.

#### Recombination between Pasteurellaceae species

Uptake of DNA from related species can also influence recombination, either directly if the DNA is sufficiently similar to recombine with the *H. influenzae* genome, or indirectly if it successfully competes with *H. influenzae* DNA for uptake or, once inside the cell, for access to nucleases or recombination machinery. For *H. influenzae* the most important competition will be with other Pasteurellaceae that share both the respiratory tract niche and the *Hin*-type USS, but similar effects are expected for Pasteurellaceae in other host species.

#### Competition with DNAs from human cells and other respiratory bacteria

In the respiratory tract, the most important source of competing DNA is human cells. However, our analysis suggests that *H. influenzae*’s uptake specificity allows its DNA to outcompete human and other foreign DNAs even if these are in 100-fold excess, a combined effect of the low number of USS_10_s and their poor match to the uptake motif. Note that efficient self-uptake does not necessarily imply a selective advantage, since USS accumulation in *H. influenzae*’s genome may simply be due to the molecular drive process.

Uptake of DNA in the respiratory track could also be influenced by the presence of chromatin and nucleoid proteins stably bound to the DNA. Although laboratory experiments typically use highly purified DNA, DNA released by cell death will be coated with these proteins, which can contribute significantly to biofilm stability (Brockman et al., 2018). Because such proteins could interfere with uptake both directly, by blocking binding to the USS, and indirectly, by blocking sliding of non-specifically bound uptake machinery along the DNA, it will be important to re-examine DNA uptake using DNA that retains its bound proteins.

## LIMITATIONS OF THE STUDY

Three factors limited measurements of uptake ratios: low sequencing read coverage, contamination of recovered donor DNA with recipient DNA, and segments of sequence identity between donor and recipient. Many reads had to be excluded at the contamination-correction step, because they were in segments that were either identical between donor and recipient or were repeated within the donor genome. Strong sequence-dependent variation in read coverage caused other positions to be excluded from analysis because they had no coverage in the control input sample. Overall, 2.3% of the genome was excluded from analysis, and an additional 1.7% was flagged as unreliable due to low coverage. In addition, the stochastic variation at low coverage positions introduced substantial noise into the calculation of experimental uptake ratios, especially at uptake valleys. On the other hand, the model predictions for long-fragment may be more accurate than indicated by their modest correlation with the noisy measured uptake ratios.

## RESOURCE AVAILABILITY

### Lead Contact

Further information and requests should be directed to and will be fulfilled by the Lead Contact. Rosemary J. Redfield

## Materials Availability

All bacterial strains are available from Joshua Chang Mell.

## Data and Code Availability

All fastQ files have been deposited at NCBI under BioProject PRJNA387591. The corresponding BioSamples are listed in Table S2. FastQ files were also deposited at Mendeley data (DOI:10.17632/hcxp9d4zkf.1. Available at https://data.mendeley.com/datasets/hcxp9d4zkf/1). The PacBio-sequenced PittGG genome reference was deposited into Genbank under SRA number SRR10207558. Full calculations, and R scripts are available at: https://github.com/mamora/DNA_uptake.

## ACKNOWLEDGEMENTS

Financial support for this work was provided by the National Science and Engineering Research Council of Canada (to RJR) and by the National Institutes of Health (to GDE (5R01DC002148-21)). We thank the Drexel Genomic Core Facility for DNA sequencing, Rachel Simister for her assistance with the bioanalyzer fragment analysis, and Matt Pennell, Sally Otto, Stephen Hallam and David Baltrus for advice on the manuscript. We also thank two anonymous reviewers for their very helpful comments, and the Editor for their patience during the revisions.

## AUTHOR CONTRIBUTIONS

MM, JCM and RJR designed the experiments and analysis. MM performed the experiments and most of the analyses; JCM and RJR performed the remaining analyses. Sequencing was done in the laboratory of GDE; data curation by RLE, JCM and MM. RJR and MM wrote the manuscript with input from the other authors.

