## Supplemental Material for "Genome-wide analysis of DNA uptake across the outer membrane of naturally competent *Haemophilus influenzae*"

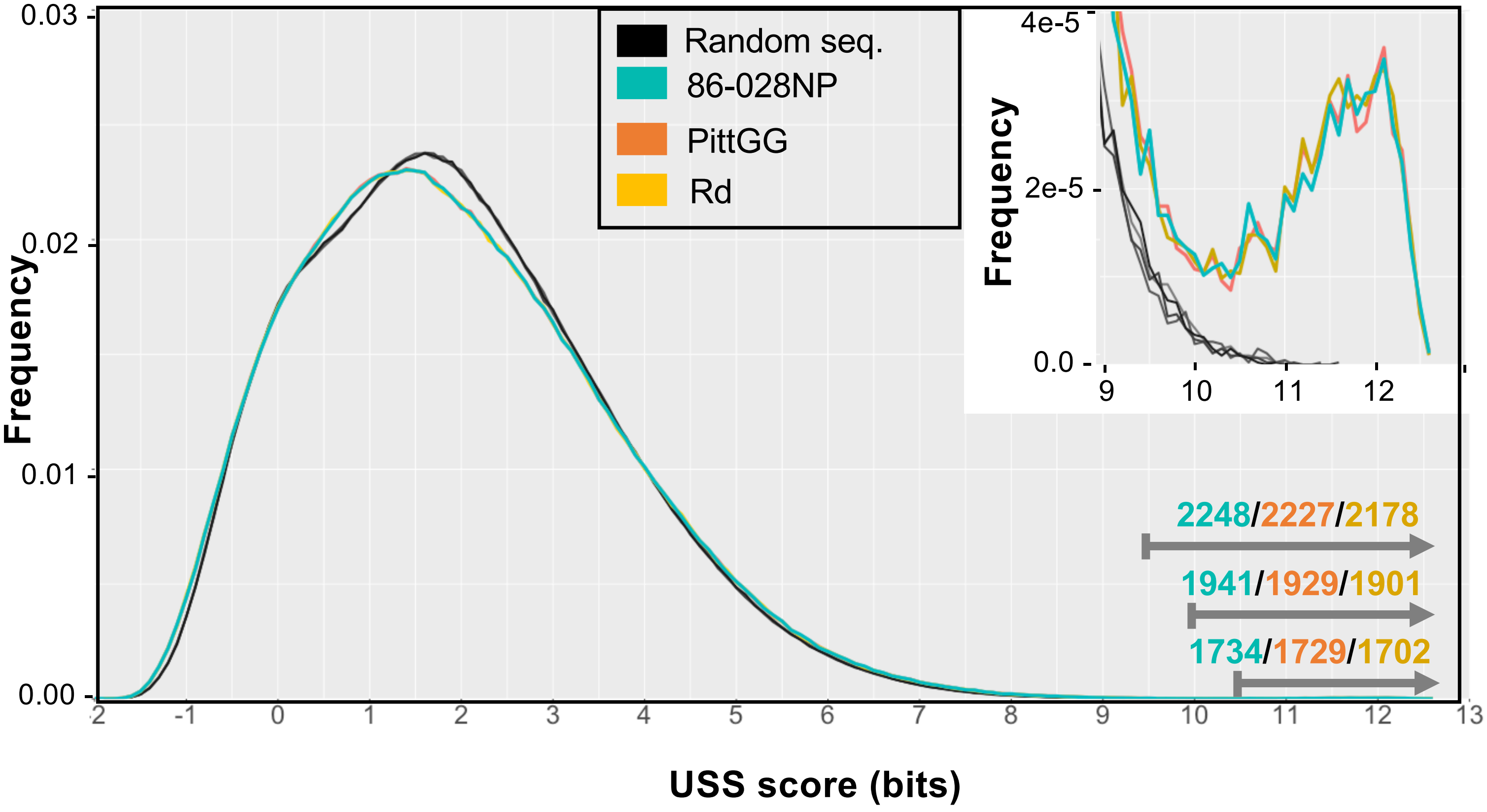

**Figure S1.** Frequency distribution of USS scores for all positions in *H. influenzae* and random-sequence genomes. Related to Figure 1.

**Legend:** 86-028 NP (blue), PittGG (orange), Rd (gold), and four random-sequence 1.9 Mb genomes with the same base composition (38%G+C, black and grey). Scores were calculated with the uptake scoring matrix in Table S1. The numbers in the lower right are the numbers of positions meeting cutoff scores of 9.5, 10.0 and 10.5 bits. **Inset:** Expanded view for positions with scores higher than 9 bits.

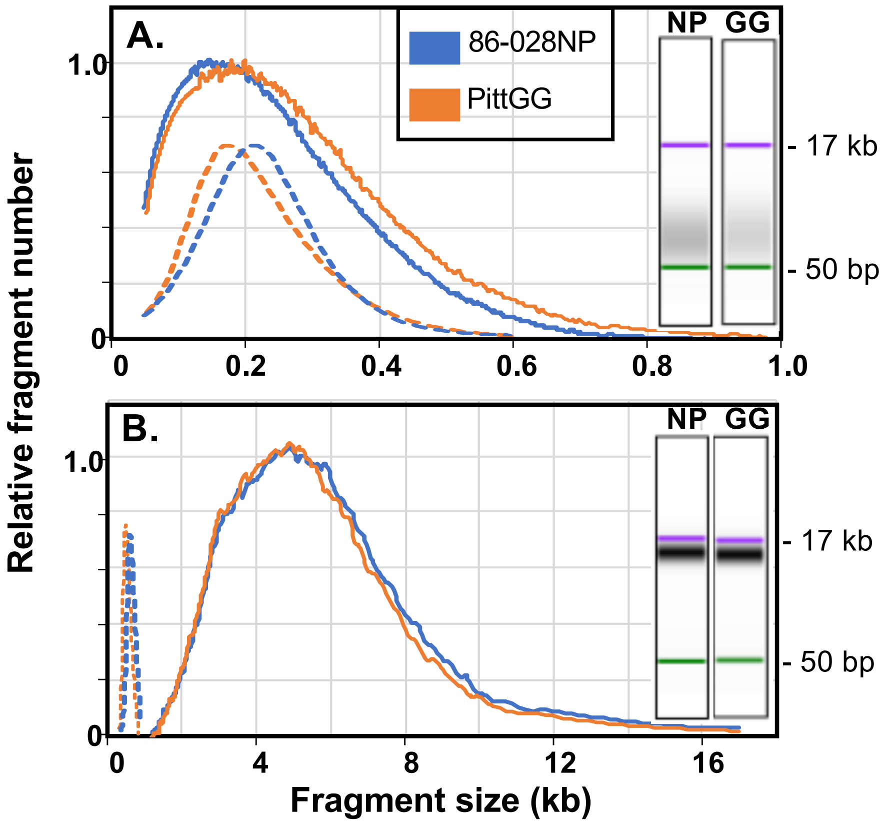

**Figure S2:** Distributions of fragment lengths in input DNA preparations. Related to Figures 3, 5 and 9.

**Legend:** Relative abundances of DNA fragment lengths were estimated from Bioanalyzer data for input DNA samples from strains 86-028NP (blue) and PittGG (orange). **A**. Short-fragment preparations. **B**. Long-fragment preparations. Solid lines: length distributions of fragments in input DNA preparations, normalized to most frequent length. Dashed lines: Length distributions of sequenced fragments, with arbitrary scaling. Insets: Bioanalyzer pseudo-gel images of sheared DNAs (NP: 86-028NP, GG: PittGG). Bioanalyzer molecular weight markers are shown in purple and green.

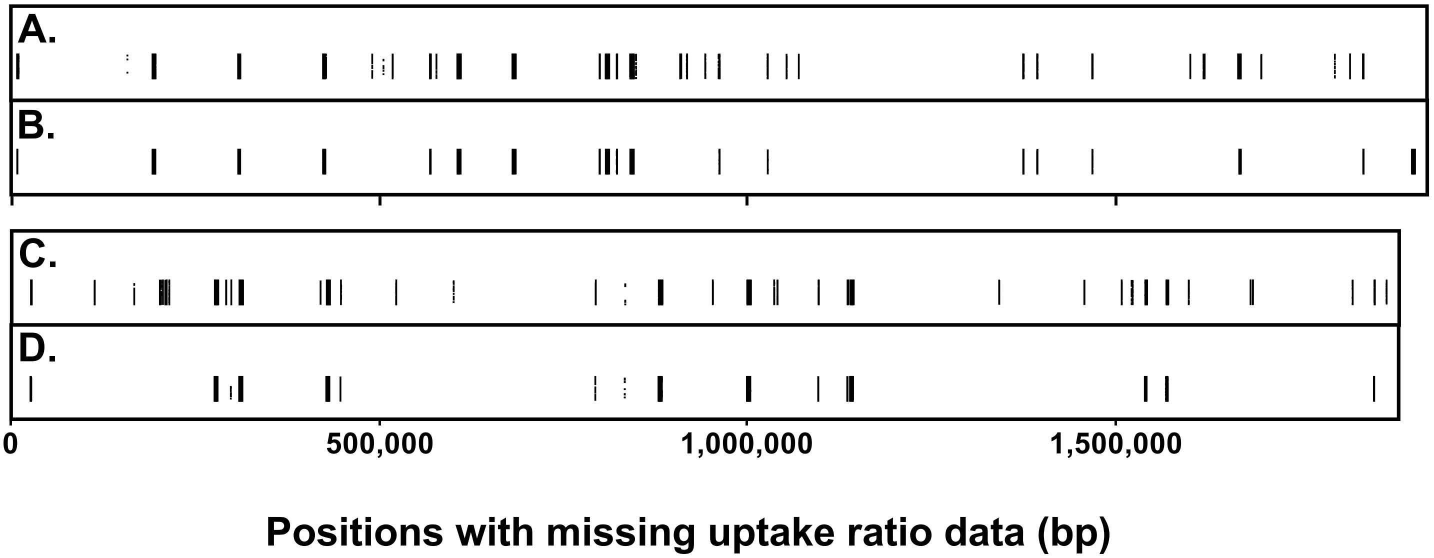

**Figure S3** Locations of positions with missing uptake ratio data. Related to Figures 3, 5 and 9.

**Legend:** Each point represents a genome position for which an uptake ratio could not be calculated. The points are vertically jittered, so segments with no coverage appear as black rectangles. **A.** 86-028NP, short-fragment data. **B.** 86-028NP, long-fragment data. **C.** PittGG, short-fragment data. **D.** PittGG, long-fragment data.

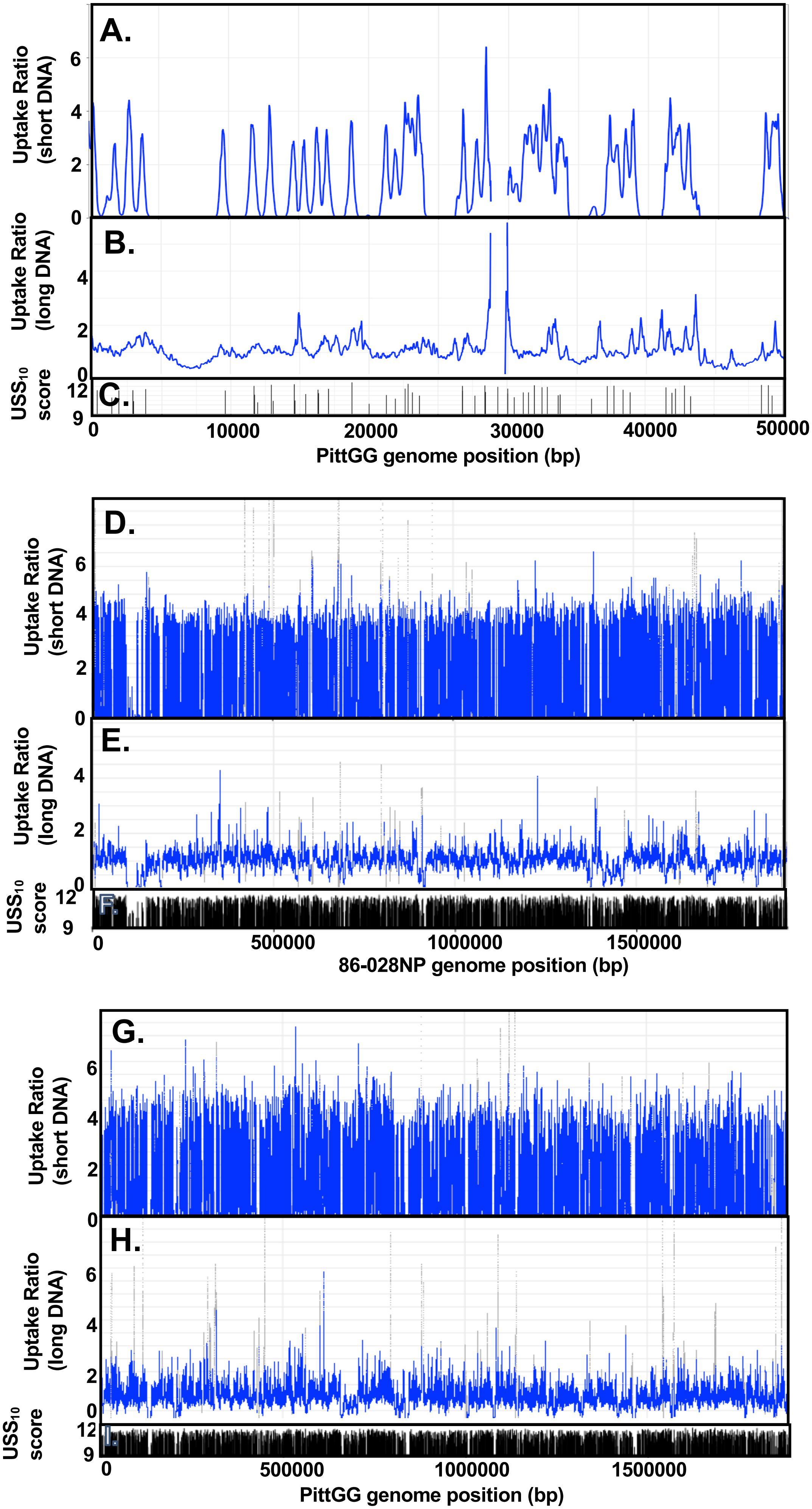
A.
Uptake Ratio (short DNA)
B.
Uptake Ratio (long DNA)
C.
USS10 score
PittGG genome position (bp)
D.
Uptake Ratio (short DNA)
E.
Uptake Ratio (long DNA)
F.
USS10 score
86-028NP genome position (bp)
G.
Uptake Ratio (short DNA)
H.
Uptake Ratio (long DNA)
I.
USS10 score
PittGG genome position (bp)

**Figure S4.** Experimentally determined uptake ratio maps and $USS_{10}$ maps. Related to Figure 3.

**Legend:** Grey points indicate positions with input coverage lower than 20 reads. Gaps indicate unmappable positions. **A-C:** Maps of a 50 kb segment of the PittGG genome. **A.** Uptake ratios of short-fragment PittGG DNA. **B.** Uptake ratios of long-fragment PittGG DNA. **C.** PittGG $USS_{10}$ positions and scores. **D-I**: Whole-genome maps. **D.** Uptake ratios of short-fragment 86-028NP DNA. **E.** Uptake ratios of long-fragment 86-028NP DNA. **F.** 86-028NP $USS_{10}$ positions and scores. **G.** Uptake ratios of short-fragment PittGG DNA. **H.** Uptake ratios of long-fragment PittGG DNA. **I.** PittGG $USS_{10}$ positions and scores.

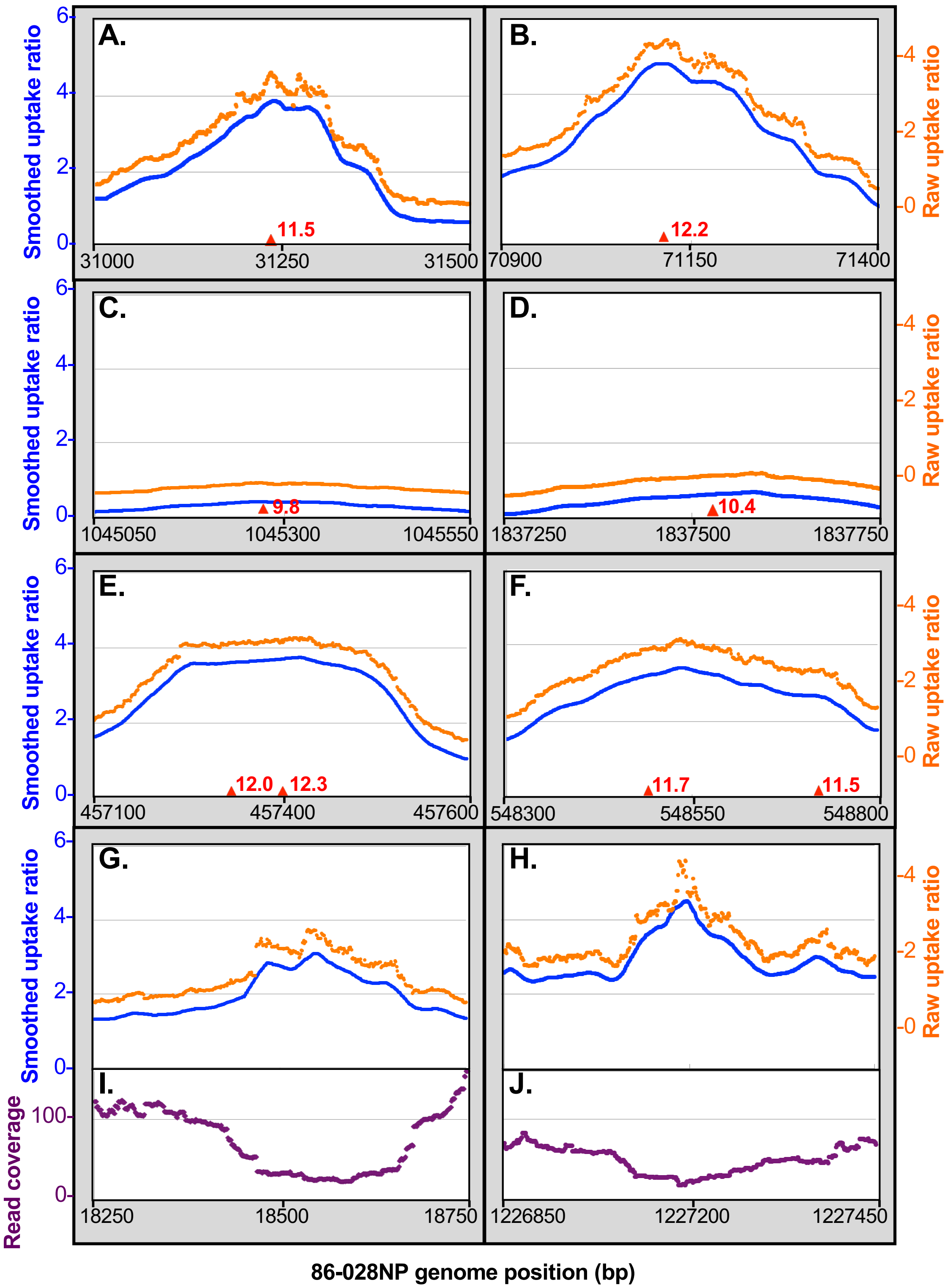
A.
B.
C.
D.
E.
F.
G.
H.
I.
J.
Smoothed uptake ratio
Raw uptake ratio
Read coverage
11.5
12.2
9.8
10.4
12.0
12.3
11.7
11.5
31000
31250
31500
70900
71150
71400
1045050
1045300
1045550
1837250
1837500
1837750
457100
457400
457600
548300
548550
548800
18250
18500
18750
1226850
1227200
1227450
86-028NP genome position (bp)

**Figure S5.** Shapes of typical 86-028NP uptake peaks. Related to Figures 3 and 9.

**Legend.** Blue dots: uptake ratios after smoothing with a 31 bp window. **A.-H.**: Orange dots: uptake ratios without smoothing (note that Y axis is offset by 0.5 units). Red triangles and numbers: locations and scores of USS. **A.-F**. 86-028NP short-fragment DNA: **A.** and **B.**: peaks at strong USS. **C.** and **D.**: peaks at weak USS. **E.** and **F.**: peaks at pairs of USS separated by **A.** 69 bp and **B.** 230?? bp. **G.** and **H.** Uptake ratio spikes not at USS in 86-028NP long-fragment DNA. **I.** and **J.** Purple dots: sequencing coverage of input 86-028NP long-fragment DNA.

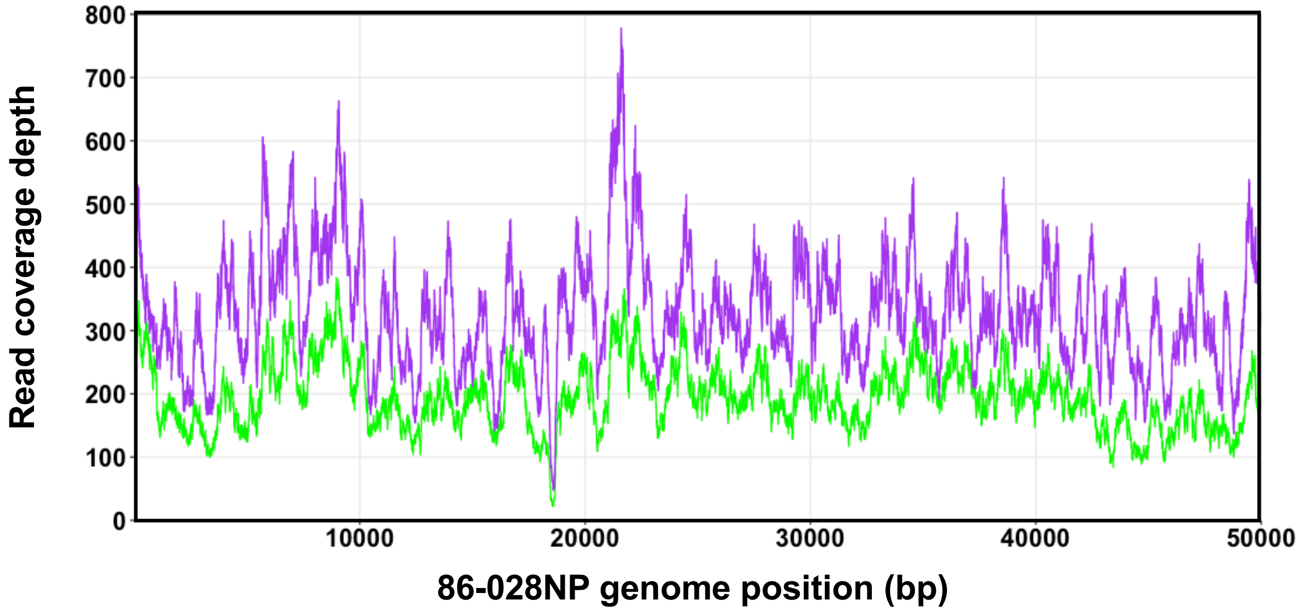

**Figure S6.** Variation in read coverage. Related to Figures 3 and 9.

**Legend:** Read coverage of the 86-028NP long-fragment (green) and short-fragment (purple) input samples over a 50 kb genome segment.

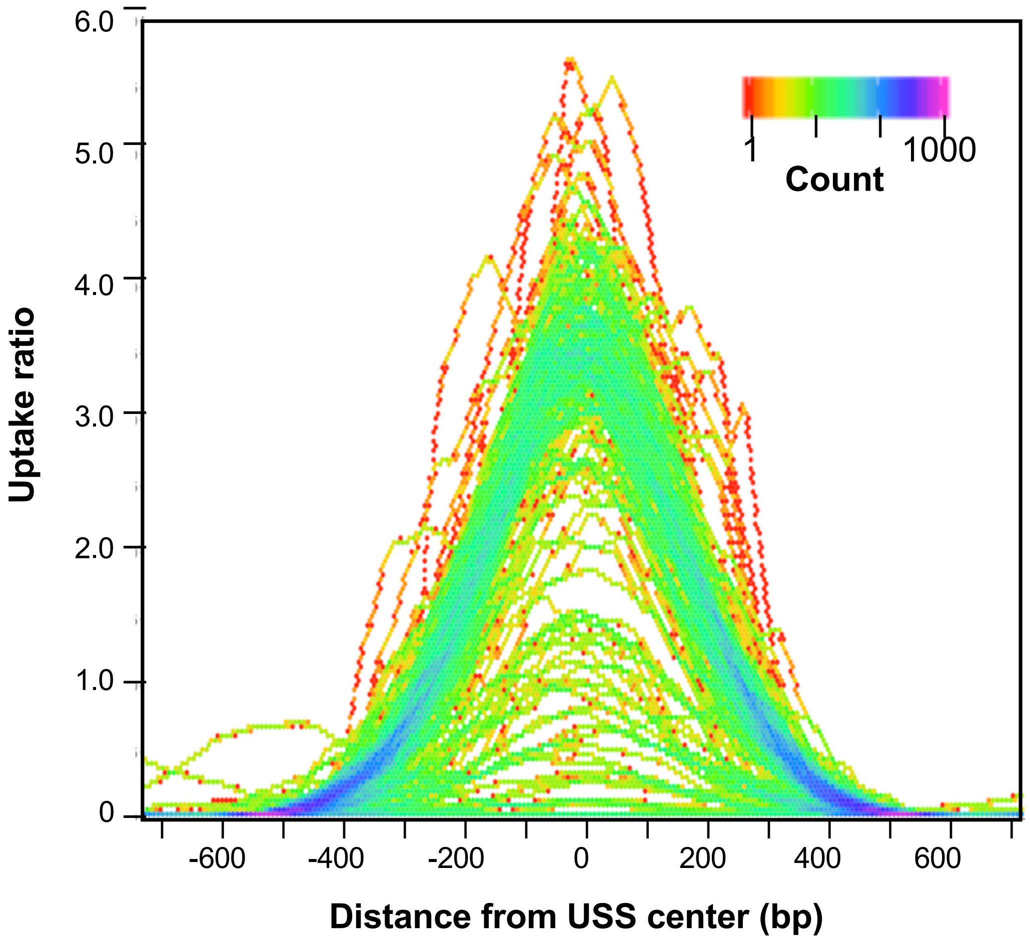

**Figure S7.** Shape analysis of isolated $USS_{10}$ peaks. Related to Figures 4 and 6.

**Legend:** Short-fragment uptake ratio data for positions around 158 86-028NP $USS_{10}$s that were separated by at least 1200 bp from other $USS_{10}$s and had uptake ratios of at least 3.0.

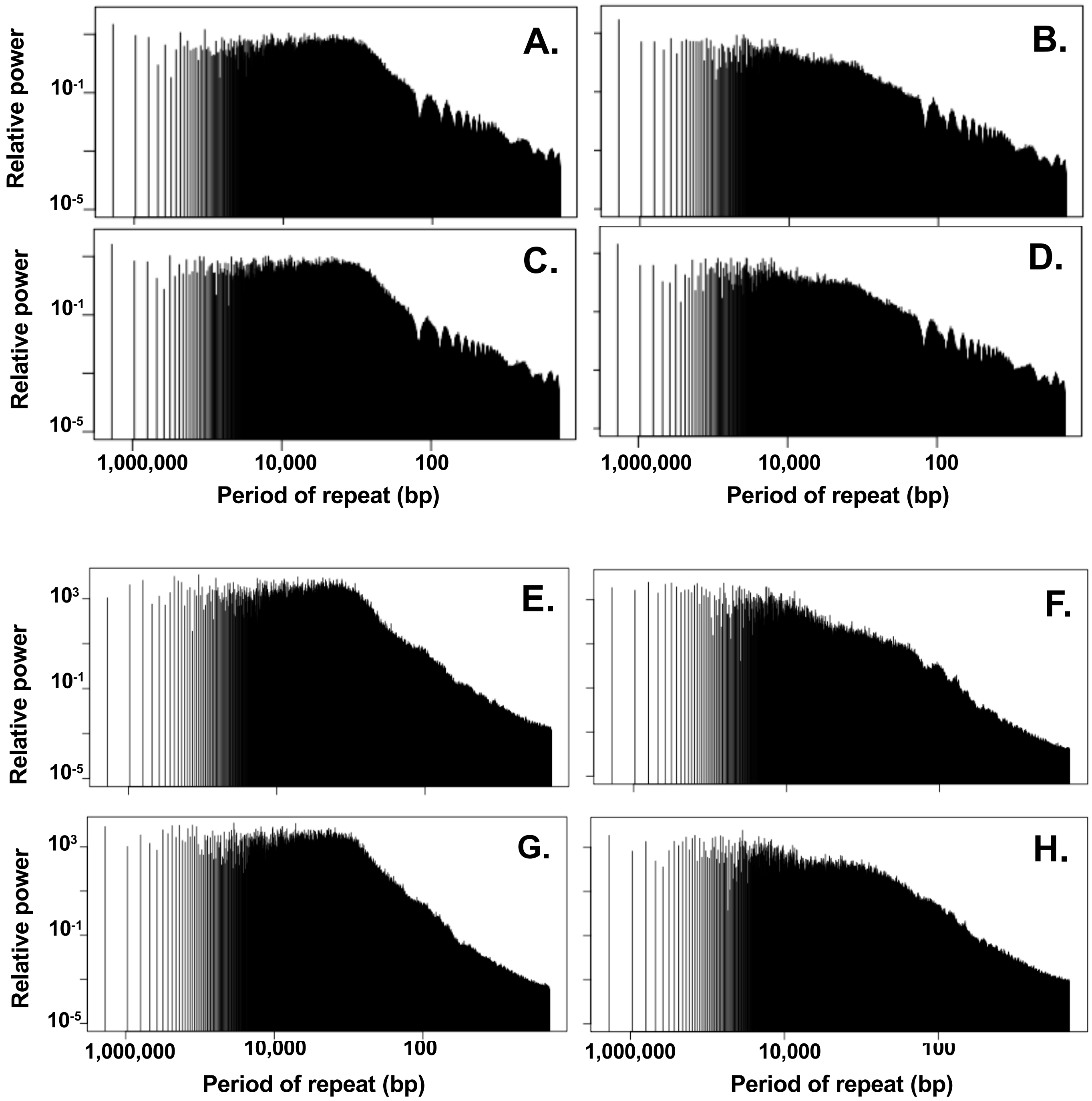

**Figure S8.** Tests of periodicity. Related to Figures 3 and 9.

**Legend:** Fourier-transform analyses were performed using R-package RCA. The X-axes are $\log_{10}$ of repeat period in bp; Y-axes are $\log_{10}$ of the relative periodicity at each repeat period. **A-D.** Tests using coverage in input samples. **E-H.** Tests using uptake ratios. Samples: **A** & **E**: 86-028NP short fragments; **B** & **F**: 86-028NP long-fragments; **C** & **G**:Pitt GG short fragments; **D** & **H**: PittGG long fragments.

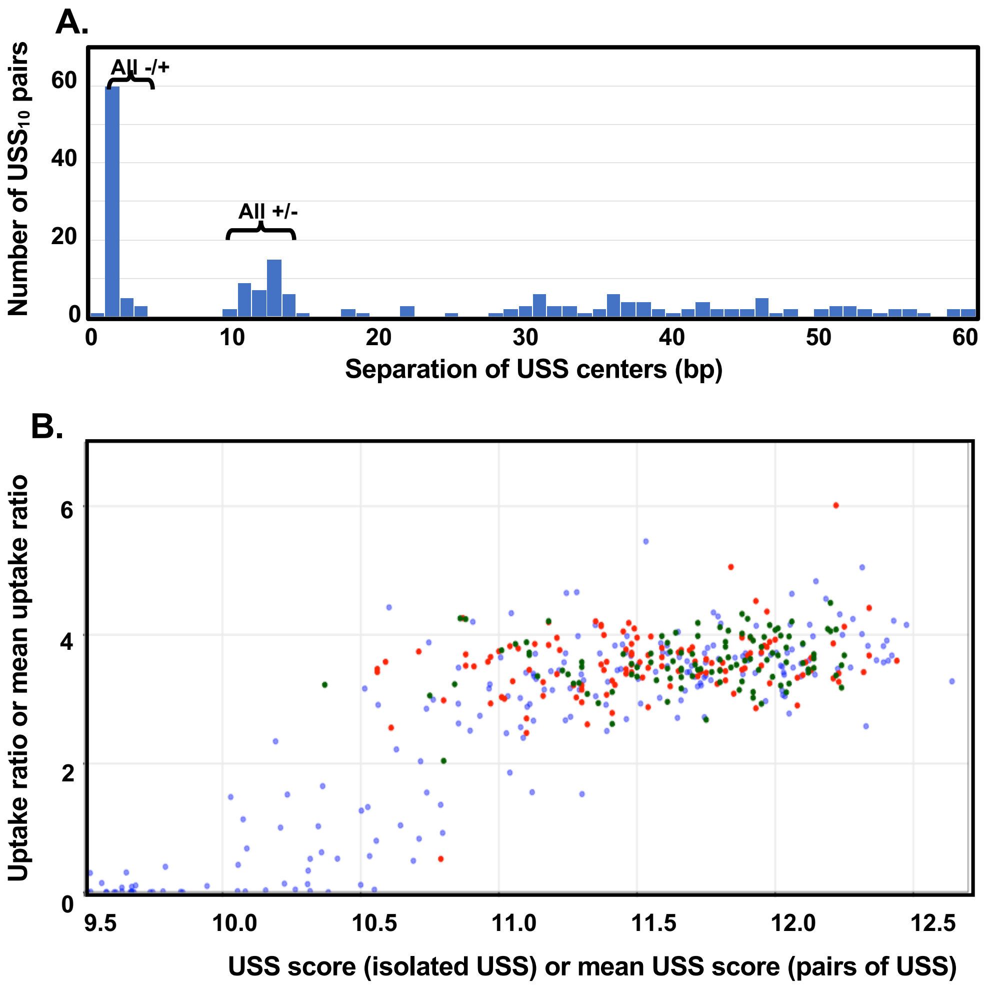

**Figure S9.** Analysis of DNA uptake effects of $USS_{10}$ pairs in the 86-028NP genome. Related to Figure 3 and 6.

**Legend: A.** Frequencies of spacings between close $USS_{10}$ pairs. **B.** Uptake ratios for isolated $USS_{10}$ (blue points, data from Figure 6) and for centers of pairs of $USS_{10}$ whose centers are 14-100 bp apart (red points) or 0-13 bp apart (dark green points).

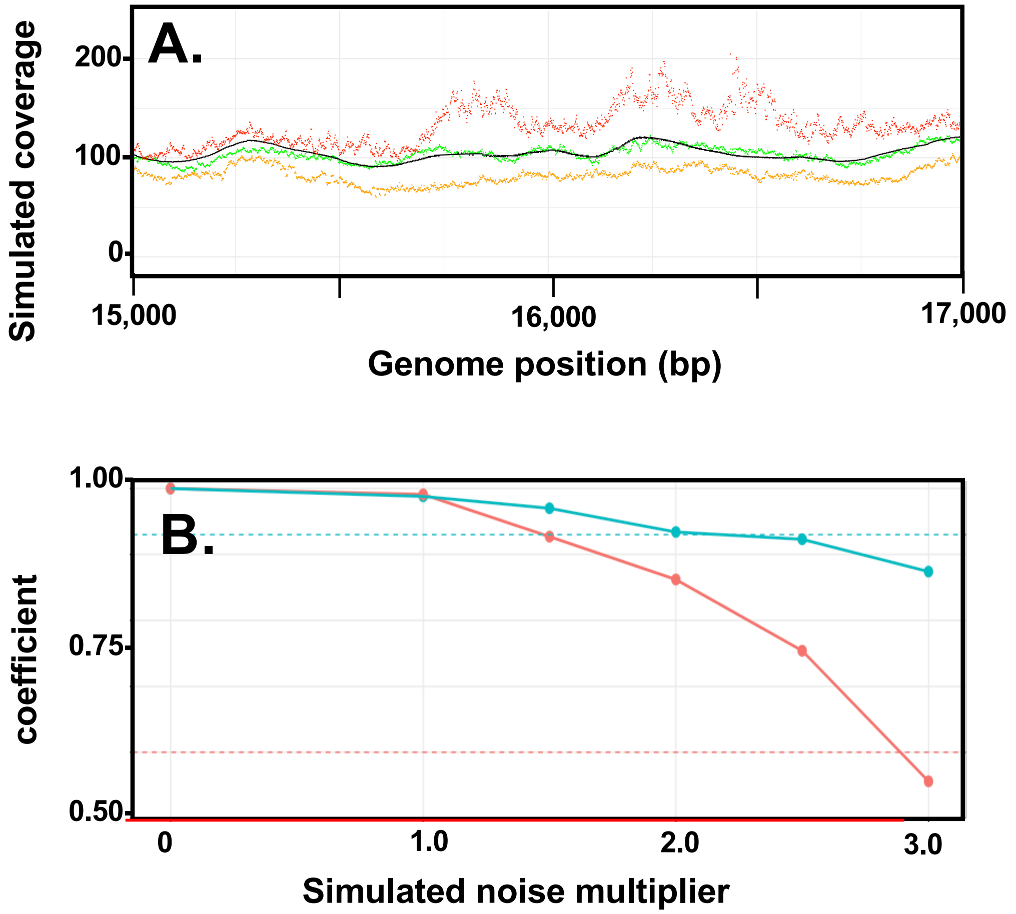

**Figure S10**. Simulated noise analysis. Related to Figures 5 and 9.

**Legend: A.** Effects of added red noise on simulated noise-free coverage. Black points: no added noise; Green, yellow and red points: simulated noise added with multipliers of 1.0, 2.0 and 3.0 respectively. **B**. Correlation coefficient from simulated uptake ratios with and without different levels of noise. X-axis represents the multiplicative factor applied to the coverage-dependent amount of noise added. Simulated results for 86-028NP-short and 86-028NP-long DNA fragments are shown in blue and red, respectively. Dashed lines at 0.93 (blue) and 0.60 (red) indicate the real-data correlation of predicted uptake with observed uptake ratios for 86-028NP-short (top) and long (bottom) fragment size distributions, respectively.

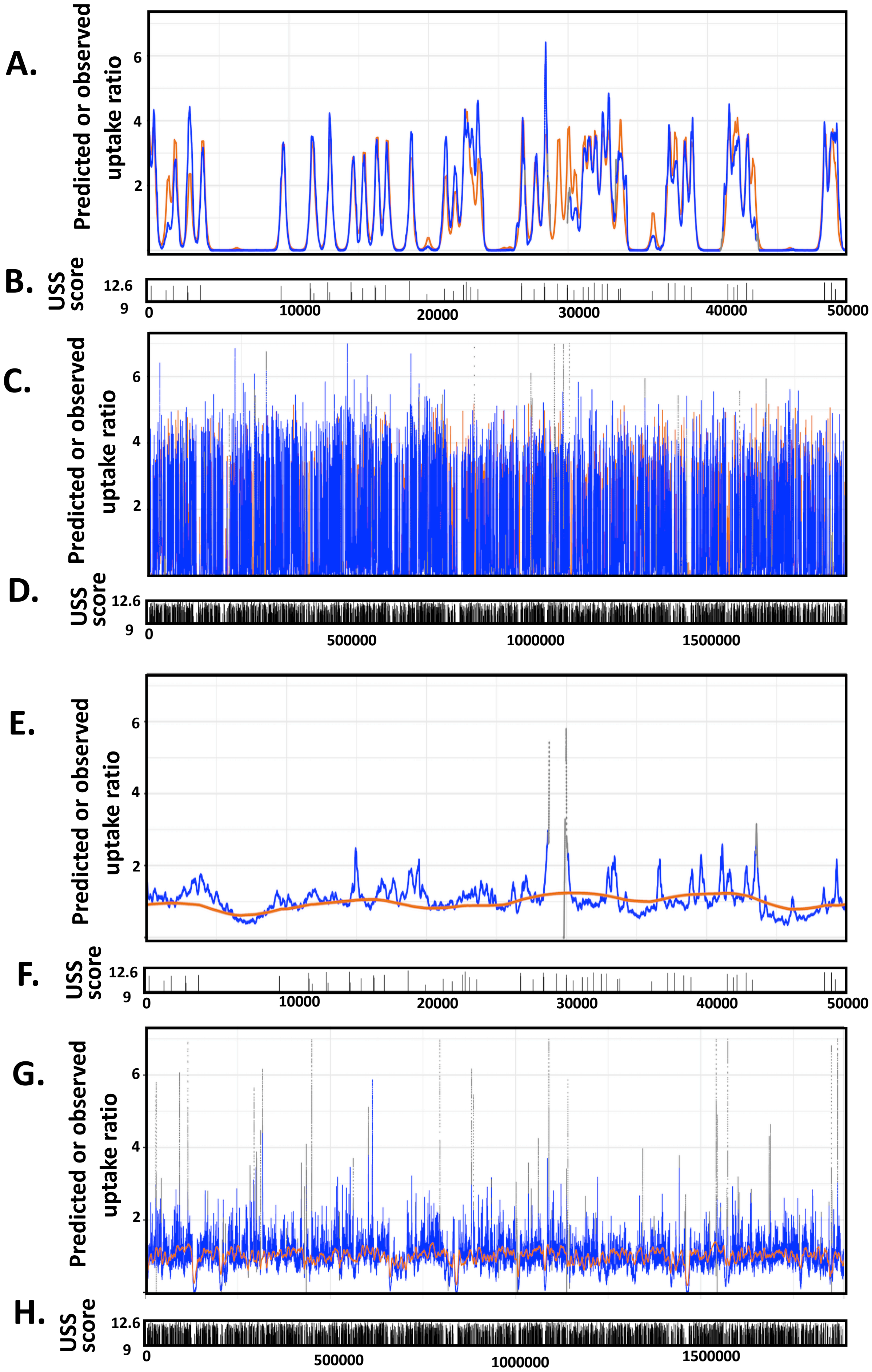
A.
Predicted or observed uptake ratio
B.
USS score
12.6
9
0
10000
20000
30000
40000
50000
C.
Predicted or observed uptake ratio
D.
USS score
500000
1000000
1500000
E.
Predicted or observed uptake ratio
F.
USS score
G.
Predicted or observed uptake ratio
H.
USS score

**Figure S11.** Predicted and observed uptake ratios for PittGG short and long DNA fragments. Related to Figure 3.

**Legend: A** and **E**: Uptake maps for the first 50kb of the PittGG genome; **C.** and **G.**: Uptake maps for the full PittGG genome. **B.**, **D.**, **F.** and **I.**: Vertical tick marks indicate locations and scores of $USS_{10}$s. Orange lines: USS-dependent uptake predicted by the revised model. Blue lines: Mean uptake ratios from 3 replicate experiments (grey points indicate positions with <20 reads input coverage, gaps indicate unmapable possitions). Note that some grey dots are beyond the tops of some panels.

**Table S1: USS uptake scoring matrix: Related to Figure 1.**

| | 1 | 2 | 3 | 4 | 5 | 6 | 7 | 8 | 9 | 10 |
|---|---|---|---|---|---|---|---|---|---|---|
| A | 0.07 | 0.09 | 0.42 | 0.63 | -0.13 | -0.13 | -0.07 | -0.06 | -0.07 | -0.10 |
| C | -0.09 | -0.04 | -0.13 | -0.12 | -0.11 | -0.12 | -0.10 | 1.86 | -0.09 | -0.12 |
| G | 0.06 | -0.02 | -0.03 | -0.07 | 1.01 | -0.13 | 1.61 | -0.05 | 1.76 | 1.57 |
| T | -0.01 | -0.03 | -0.08 | -0.06 | -0.12 | 0.83 | -0.12 | -0.05 | -0.08 | -0.11 |

| | 11 | 12 | 13 | 14 | 15 | 16 | 17 | 18 | 19 | 20 |
|---|---|---|---|---|---|---|---|---|---|---|
| A | -0.11 | -0.01 | 0.15 | 0.07 | 0.04 | 0.04 | -0.02 | -0.05 | -0.02 | 0.00 |
| C | -0.03 | 0.05 | -0.07 | -0.05 | -0.08 | -0.08 | -0.04 | 0.01 | -0.02 | -0.01 |
| G | -0.10 | -0.06 | 0.03 | 0.00 | -0.06 | -0.11 | -0.11 | -0.08 | -0.02 | 0.00 |
| T | 0.40 | 0.03 | -0.07 | -0.02 | 0.13 | 0.27 | 0.26 | 0.16 | 0.07 | 0.00 |

| | 21 | 22 | 23 | 24 | 25 | 26 | 27 | 28 | 29 | 30 | 31 |
|---|---|---|---|---|---|---|---|---|---|---|---|
| A | 0.00 | 0.00 | -0.01 | 0.00 | 0.05 | -0.01 | 0.04 | -0.06 | -0.06 | -0.03 | -0.03 |
| C | 0.01 | 0.00 | 0.00 | 0.01 | -0.06 | -0.04 | -0.10 | -0.01 | -0.03 | -0.01 | -0.01 |
| G | 0.01 | 0.01 | 0.01 | 0.00 | 0.06 | -0.06 | -0.13 | -0.12 | -0.08 | -0.03 | 0.00 |
| T | -0.02 | -0.01 | 0.00 | -0.01 | -0.04 | 0.13 | 0.35 | 0.32 | 0.23 | 0.08 | 0.04 |

#### Table S2. Sample metadata. Related to Table 1 and Figures 3 and 9.

| Sample name[1] | Sample type | Biosample ID | Bioproject accession | Donor strain | Genome size | Fragment size range (bp) | Recipient strain | % of DNA recovered |
|---|---|---|---|---|---|---|---|---|
| **UP1 (NP lg)** | Taken-up DNA | SAMN07187224 | PRJNA387591 | 86-028NP NalR | 1,914,387 | 1500-17000 | RR3117 (*rec2::spec*) | 2.06 |
| **UP2 (NP lg)** | Taken-up DNA | SAMN07187225 | PRJNA387591 | 86-028NP NalR | 1,914,387 | 1500-17000 | RR3125 (*rec2-*) | 1.38 |
| **UP3 (NP lg)** | Taken-up DNA | SAMN07187226 | PRJNA387591 | 86-028NP NalR | 1,914,387 | 1500-17000 | RR3125 (*rec2-*) | 1.38 |
| **UP4 (GG lg)** | Taken-up DNA | SAMN07187227 | PRJNA387591 | PittGG) | 1,887,046 | 1500-17000 | RR3117 (*rec2::spec*) | 1.60 |
| **UP5 (GG lg)** | Taken-up DNA | SAMN07187228 | PRJNA387591 | PittGG | 1,887,046 | 1500-17000 | RR3125 (*rec2-*) | 1.24 |
| **UP6 (GG lg)** | Taken-up DNA | SAMN07187229 | PRJNA387591 | PittGG (RR1361) | 1,887,046 | 1500-17000 | RR3125 (*rec2-*) | 2.48 |
| **UP7 (NP sh)** | Taken-up DNA | SAMN07187230 | PRJNA387591 | 86-028NP NalR | 1,914,387 | 50-800 | RR3117 (*rec2::spec*) | 0.64 |
| **UP8 (NP sh)** | Taken-up DNA | SAMN07187231 | PRJNA387591 | 86-028NP NalR | 1,914,387 | 50-800 | RR3125 (*rec2-*) | 0.32 |
| **UP9 (NP sh)** | Taken-up DNA | SAMN07187232 | PRJNA387591 | 86-028NP NalR | 1,914,387 | 50-800 | RR3125 (*rec2-*) | 0.41 |
| **UP10 (GG sh)** | Taken-up DNA | SAMN07187233 | PRJNA387591 | PittGG | 1,887,046 | 50-800 | RR3117 (*rec2::spec*) | 0.77 |
| **UP11 (GG sh)** | Taken-up DNA | SAMN07187234 | PRJNA387591 | PittGG | 1,887,046 | 50-800 | RR3125 (*rec2-*) | 0.91 |
| **UP12 (GG sh)** | Taken-up DNA | SAMN07187235 | PRJNA387591 | PittGG | 1,887,046 | 50-800 | RR3125 (*rec2-*) | 1.03 |
| **UP13 (NP lg)** | Input DNA | SAMN07187236 | PRJNA387591 | 86-028NP NalR | 1,914,387 | 1500-17000 | N/A | N/A |
| **UP14 (GG lg)** | Input DNA | SAMN07187237 | PRJNA387591 | PittGG | 1,887,046 | 1500-17000 | N/A | N/A |
| **UP15 (NP sh)** | Input DNA | SAMN07187238 | PRJNA387591 | 86-028NP NalR | 1,914,387 | 50-800 | N/A | N/A |
| **UP16 (GG sh)** | Input DNA | SAMN07187239 | PRJNA387591 | PittGG | 1,887,046 | 50-800 | N/A | N/A |
| **Rd** | recipient DNA | SAMN12049038 | PRJNA387591 | Rd (recipient) | 1,831,585 | Not sheared | Rd KW20 | N/A |

| Sample name[1] | Mean size of library fragments | Un-mapped reads | Mapped reads | Mean MAPQ score | Reads mapping only to recipient (MAPQ>0) | Reads mapping only to donor (MAPQ>0) | % contam-ination with Rd DNA | % of reads removed | Mean read coverage |
|---|---|---|---|---|---|---|---|---|---|
| **UP1 (NP lg)** | 342.4 | 22,982 | 2,6468,71 | 47.2 | 122,712 | 2,253,377 | 5.16 | 14.9 | 174 |
| **UP2 (NP lg)** | 334.0 | 29,276 | 2,386,785 | 47.1 | 151,131 | 1,987,921 | 7.07 | 16.7 | 154 |
| **UP3 (NP lg)** | 341.3 | 29,111 | 2,962,387 | 47.1 | 226,132 | 2,428,674 | 8.52 | 18.0 | 188 |
| **UP4 (GG lg)** | 348.8 | 11,134 | 2,183,783 | 46.3 | 145,426 | 1,792,255 | 7.51 | 17.9 | 140 |
| **UP5 (GG lg)** | 348.2 | 10,423 | 2,149,185 | 46.4 | 366,986 | 1,539,930 | 19.25 | 28.3 | 121 |
| **UP6 (GG lg)** | 309.5 | 7,732 | 1,227,888 | 46.0 | 97,875 | 978,918 | 9.09 | 20.3 | 77 |
| **UP7 (NP sh)** | 227.8 | 203,460 | 5,011,237 | 45.8 | 363,683 | 4,006,647 | 8.32 | 20.0 | 303 |
| **UP8 (NP sh)** | 219.5 | 211,676 | 3,592,281 | 44.7 | 522,918 | 2,591,844 | 16.79 | 27.8 | 197 |
| **UP9 (NP sh)** | 247.2 | 201,601 | 7,0983,09 | 46.5 | 801,198 | 5,452,848 | 12.81 | 23.2 | 417 |
| **UP10 (GG sh)** | 240.9 | 171,799 | 4,746,454 | 45.4 | 323,716 | 3,807,524 | 7.84 | 19.8 | 293 |
| **UP11 (GG sh)** | 241.0 | 124,946 | 5,563,501 | 46.1 | 157,046 | 4,687,429 | 3.24 | 15.7 | 361 |
| **UP12 (GG sh)** | 233.8 | 234,499 | 5,298,060 | 44.9 | 205,997 | 4,389,741 | 4.48 | 17.1 | 337 |
| **UP13 (NP lg)** | 344.5 | 11,787 | 2,715,317 | 46.8 | 1,333 | 24,04,540 | 0.06 | 11.4 | 185 |
| **UP14 (GG lg)** | 388.4 | 7,774 | 5,836,702 | 46.4 | 1,345 | 5,314,003 | 0.03 | 9.0 | 192 |
| **UP15 (NP sh)** | 222.4 | 13,750 | 4740911 | 45.3 | 1154 | 3,961,509 | 0.03 | 16.4 | 296 |
| **UP16 (GG sh)** | 234.0 | 9531 | 6627713 | 45.3 | 1668 | 4,938,510 | 0.03 | 25.5 | 395 |
| **Rd** | 241.0 | 4564 | 4083313 | N/A | N/A | N/A | N/A | N/A | 298 |

1. **NP lg:** Long-fragment 86-028NP DNA, **GG lg:** Long-fragment PittGG DNA, **NP-sh:** Short-fragment 86-028NP DNA, **GG sh:** Short-fragment PittGG DNA.

#### Table S3. Analysis of low-coverage positions. Related to Figures 3 and 9.

| 86-028NP-long: | | | | | | | |
|---|---|---|---|---|---|---|---|
| **Input coverage:** | **≤10** | **10-20** | **20-50** | **50-100** | **>100** | **sum** | **% in the genome** |
| **% of positions with this coverage** | 2.36 | 0.45 | 1.78 | 6.14 | 89.27 | 100 | |
| **% of positions with uptake ratios >2.0** | 7.53 | 4.73 | 20.75 | 37.81 | 29.18 | 100 | 0.82% |
| **% of positions with uptake ratios >3.0** | 30.22 | 12.07 | 38.38 | 19.33 | 0 | 100 | 0.06% |
| **% of positions with uptake ratios <0.5** | 1.78 | 0.85 | 2.7 | 6.6 | 88.07 | 100 | 8.90% |
| **% of positions with uptake ratios <0.25** | 2.19 | 0.69 | 2.08 | 4.82 | 90.22 | 100 | 4.00% |
| **86-028NP-short** | | | | | | | |
| **Input coverage** | **≤10** | **10-20** | **20-50** | **50-100** | **>100** | **sum** | |
| **% of positions with this coverage** | 3.37 | 0.63 | 1.68 | 2.88 | 91.44 | 100 | |
| **% of positions with uptake ratios >4.0** | 9.13 | 3.93 | 7.11 | 9.2 | 70.63 | 100 | 1.11% |
| **% of positions with uptake ratios >5.0** | 45.46 | 16.48 | 7.87 | 9.11 | 21.08 | 100 | 0.13% |
| **% of positions with uptake ratios >0.1** | 1.19 | 0.75 | 1.91 | 3.23 | 92.92 | 100 | 44.40% |
| **% of positions with uptake ratios >0.01** | 1.45 | 0.76 | 1.93 | 3.22 | 92.63 | 100 | 28.15% |
| **PittGG-long** | | | | | | | |
| **Input coverage** | **≤10** | **10-20** | **20-50** | **50-100** | **>100** | **sum** | |
| **% of positions with this coverage** | 2.33 | 0.43 | 1.36 | 2.85 | 93.03 | 100 | |
| **% of positions with uptake ratios >2.0** | 16.85 | 10.76 | 25.65 | 27.8 | 18.94 | 100 | 1.57% |
| **% of positions with uptake ratios >3.0** | 61.87 | 17.07 | 15.62 | 5.44 | 0 | 100 | 0.26% |
| **% of positions with uptake ratios <0.5** | 0.58 | 0.32 | 0.97 | 2.87 | 95.26 | 100 | 6.90% |
| **% of positions with uptake ratios <0.25** | 0.86 | 0.46 | 1.65 | 4.41 | 92.62 | 100 | 2.57% |
| **PttGG-short** | | | | | | | |
| **Input coverage** | **≤10** | **10-20** | **20-50** | **50-100** | **>100** | **sum** | |
| **% of positions with this coverage** | 3.11 | 0.73 | 1.98 | 2.9 | 91.28 | 100 | |
| **% of positions with uptake ratios >4.0** | 1.73 | 1.62 | 6.77 | 11.33 | 78.55 | 100 | 2.50% |
| **% of positions with uptake ratios >5.0** | 8.8 | 7.41 | 20.2 | 28.53 | 35.06 | 100 | 0.24% |
| **% of positions with uptake ratios >0.1** | 1.25 | 0.77 | 2.25 | 3.02 | 92.72 | 100 | 39.38% |
| **% of positions with uptake ratios >0.01** | 1.54 | 0.76 | 2.06 | 3.02 | 92.62 | 100 | 26.50% |

### Transparent Methods

**Identifying USSs in the genomes.** Genomic USSs were identified by scoring each genome position with the position-specific scoring matrix (PSSM) of Mell et al. (2012); this is based on uptake of synthetic fragments containing degenerate USS sequences. Positions scoring ≥ 10.0 or ≥ 9.5 (maximum score is 12.6) were included in the standard ($USS_{10}$ and $USS_{9.5}$) lists of USS locations. Since USS are asymmetric, USS positions in both orientations were specified by the location of their central base 16. Sequence logos of USSs were generated using R package seqLogo v. 3.8.

**Predicting DNA uptake from DNA sequence.** The predictive model was written in R v.3.5.1. Given a list of USS positions and scores in a DNA genome of specified length, it used a specified distribution of DNA fragment lengths or length bins (e.g. 1-100 bp, 101-200 bp, etc.) to calculate the relative uptake of every position in a circular genome. At each DNA position in turn, for each fragment length or bin, the model summed the predicted uptake contributions for every fragment of that length that overlapped the position. For efficiency, the full calculation was only done for the first position. At each subsequent position, the model calculated the new sum from the previous position's sum by subtracting the contribution of the formerly leftmost fragment and adding the contribution of the new rightmost fragment (Figure 2.3A).

Each fragment's predicted contribution to uptake depended on the number of USS it contained, and on the scores and relative locations of these USS. Fragments with no or incomplete USSs were assigned baseline values for the probabilities of being bound (*p_bind*) and taken up (*p_uptake*); initial values for both were arbitrarily set to 0.1. For fragments with one or more complete USS$_{10}$, *p_bind* was calculated as *1 – mean_gap/20000,* where *mean_gap* was the mean length of USS-free segments in the fragment and *20000* the maximum fragment length in bp. The uptake function *p_uptake* was initially specified as *p_uptake = 0.1 + (1 - 0.1)/(1 + exp(-5 * (score – 11)))*.

Once the model had calculated the contributions of a specific fragment length or length bin to uptake of every genome position, it moved on to the next length or bin. Once the contributions of every length or bin had been calculated, the model combined all the contributions for each position, taking into account the frequency of each length or bin in the input DNA. These position-specific uptake predictions were then normalized to a mean genome-wide uptake value of 1.0.

In response to ongoing analysis of the 86-028NP DNA uptake data, the initial model underwent modifications to improve its predictions for a 'far from USS$_{10}$' subset of positions that were at least 0.5kb from a USS$_{9.5.}$ (n = 361965 positions), combined with 209 USS$_{9.5}$ peak positions separated from the nearest USS$_{10}$s by at least 1000 bp. This reduced the baseline *p_uptake* of USS-free fragments from 0.1 to 0.005 and the USS cutoff score from 10 to 9.5, excluded from consideration USS$_{9.5}$ that were within 50 bp of fragment ends, and identified better slope and inflection point values for the sigmoidal uptake function using the R function "nls" from the stats-package. These changes replaced the previous uptake function with *p_uptake = 0.005 + (1 - 0.005)/(1 + exp(-3.8 * (score – 10.6)))*.

A subsequent change adjusted uptake predictions according to the GC content around each position. First the observed effect of GC content on uptake was approximated by a linear function describing how the 86-028NP long fragment uptake ratios depended on their local GC contents calculated with a 2 kb window (see inset in Fig. GC). Each genomic position was assigned the GC content of a 1001 bp window centered on it. The predicted uptake at each position was then modified using the local GC content and the function.

**Bacterial strains, culturing, and competent cell preparations:** The KW20 recipient strains were *rec2* derivatives of the standard *H. influenzae* lab strain Rd KW20, with (RR3117) and without (RR3125) a spectinomycin resistance allele (Mell et al., 2012; Sinha et al., 2012). The 86-028NP donor strain (RR3133) was a derivative with a nalidixic acid resistance allele (Mell *et al.* 2011); the PittGG isolate was unmodified. Standard growth and culturing methods were used (Poje and Redfield, 2003); liquid cultures were grown with shaking at 37 °C in brain-heart infusion broth supplemented with NAD (2µg/ml) and hemin (10 µg/ml) (sBHI), with 1.2% agar added for plate cultures. To prepare naturally competent cells, cultures were first maintained in exponential growth at $OD_{600}$ below 0.2 for at least 2 hr, and at $OD_{600} = 0.2$ cells were collected by filtration from 10 ml of culture, transferred into 10 ml of starvation medium M-IV, and incubated at 37 °C for 100 minutes before DNA uptake experiments (Poje and Redfield, 2003).

**Input DNA preparations.** High molecular weight donor DNA was purified using standard phenol:chloroform extractions (Sambrook, 2001) from 10 ml overnight cultures of the 86-028NP derivative, and PittGG carrying selectable markers (Table S2). This DNA was then sheared into separate ‘long fragment’ (1.5-9 kb) and ‘short fragment’ (50-500 bp) preparations using Covaris G-tubes and sonication respectively. The fragment length distributions were measured using a Bioanalyzer with a DNA 12000 kit (Agilent), dividing the relative fluorescence at each time point by its fragment length estimated from the size standards.

**DNA uptake and recovery.** 10 ml of competent *rec-2* mutant Rd cells in MIV were incubated with 10 µg of sheared donor DNA for 20 min at 37 °C. To degrade remaining free DNA, the culture was incubated with 1 ug/ml of DNase I for 5 minutes. Cells were washed twice by pelleting and resuspension in cold MIV, and the final pellet was rinsed twice with cold MIV before resuspension in 0.5 ml of extraction buffer (Tris-HCL 10 mM pH 7.5, EDTA 10 mM, CsCl 1.0 M). Periplasmic DNA was extracted using the organic phenol:acetone extraction method as described by Mell *et al.* 2012 (Barouki and Smith, 1985; Kahn et al., 1983; Mell et al., 2012) followed by an ethanol precipitation. DNA was resuspended in 20 µl of $T_{10}E_{10}$ buffer (Tris-HCl 10 mM pH 7.5, EDTA 10 mM). The DNA was then incubated at 37 °C with 400 ng of RNase A for 1 hour, followed by 30 min incubation with 30 ng of proteinase K to remove RNase A. Recovered DNA was then separated from longer fragments of contaminating genomic DNA by electrophoresis in a 0.8% agarose gel and recovered from the gel slice with a Zymo gel DNA recovery kit. Recovered periplasmic DNA was quantified using a Qubit dsDNA HS Assay Kit (absolute DNA concentration).

**DNA sequencing and data processing.** Sequencing libraries of the input and taken-up DNA samples were prepared using Illumina Nextera XT DNA library prep kits according to manufacturer recommendations. An Illumina NextSeq500 was used to generate 1-10 million paired-end reads of 2x150 nt for each library (giving >100-fold genomic coverage). Summary statistics for each sample are provided in Table S2.

**Reference sequences***:* The original PittGG reference (NC_009567.1) generated by pyrosequencing had many indel errors, so a new reference was constructed by Pacific Biosciences RSII of our laboratory version of this strain (RR1361) (assembly by HGAP2 v2.3, followed by Circlator (Hunt et al., 2015), and then Quiver to polish the circular junction). For our analysis the NCBI sequence references for PittGG, 86-028NP, (NC_007146.2) and Rd KW20 (NC_000907.1) were then further corrected using Pilon v1.22 and the new Illumina reads of input or control samples. This was particularly important for the Rd KW20 recipient reference, since the original sequence dates from 1995 (Fleischmann et al., 1995) and contains several hundred ambiguous bases and errors. This correction step also accommodated the presence of the nalidixic acid resistance marker in 86-028NP.

Competition essays simulations of *H. influenzae* with human DNA, used 3 random segments of the same size as *H. influenzae* 86-028NP genome (1914386 bp) the Chromosome 1 (CM000663.2, positions 33610150 – 35524535), Chromosome 3 (CM000665.2, positions 1348752 – 3263137), and Chromosome 12 (CM000674.2, positions 18170588 – 20084973). Respiratory bacterial genomes used to score $USS_{10}$ and $USS_{11}$ in table 2 and competitions essays were *Streptococcus pneumoniae* R6 (NC_003098.1), *Neisseria meningitidis* MC58 (NC_003112.2), *Pseudomonas aeruginosa* PAO1 (NC_002516.2), *Aggregatibacter actinomycetemcomitans* VT1169 (NZ_CP012958.1), *Haemophilus parainfluenzae* T3T1 (NC_015964.1), *Haemophilus ducreyi* 35000HP (NC_002940.2), *Mannheimia haemolytica* M42548 (NC_021082.1).

**Chromosomal contamination measurements and corrections:** Reads from the recipient genomic DNA that contaminated taken-up DNA samples were identified by a 'competitive alignment' step that aligned all the sample's reads (using bwa mem v0.7.15, samblaster v0.1.24, and sambamba v0.5.0) to a concatenated double-reference sequence consisting of the recipient Rd genome and the donor genome (86-028NP or PittGG). Because the donor and recipient genomes are distinguished by a high density of SNVs, as well as structural variation and large indels (Harrison et al., 2005; Hogg et al., 2007; Mell et al., 2011), most contaminating Rd reads in uptake samples aligned only to the Rd reference while most of the desired donor-derived reads aligned only to the donor reference. Reads that mapped equally well to both genomes or to repetitive sequences within a genome were identified by their MAPQ scores of 0 and were removed from the analysis. The numbers of reads mapping uniquely to either donor or recipient genome were then used to calculate the contamination level of each taken-up DNA sample, as the ratio of recipient-mapping reads to total uniquely mapping reads (Table S2). Subsequent depth of coverage values and summary statistics were extracted for all positions or specific intervals using bedtools coverage v2.16.2 or sambamba flagstat (Table S2). All subsequent analyses and plotting used the R statistical programming language, including standard add-on packages dplyr, tidyr, plyr, ggplot2, data.table. Other packages used are specified below. Code is available at https://github.com/mamora/DNA_uptake.

**Calculation of experimental uptake ratios from sequence coverage.** After contaminating reads had been removed from each sample, uptake maps for each donor DNA were created by dividing the mean of the three normalized taken-up-DNA coverages for each position by the corresponding normalized input-DNA coverage. Finally, uptake ratios were normalized to a genome-wide mean uptake of 1.0 and smoothed by calculating the mean uptake over a 31 bp central-oriented sliding window using function rollapply from R package zoo v. 1.8-5. The effects of this smoothing are shown for the peak examples in Figure S5.

**Periodicity analysis:** To detect possible periodic patterns in coverage depth and in uptake ratios for the four datasets, periodograms were created using the R package TSA v. 1.2.

**Analysis of uptake ratio data:** To obtain a set of well-isolated $USS_{10}$s for analysis of peak shapes, we identified the closest peak separation at which USS effects did not overlap by examining sets of $USS_{10}$ that were separated by different distances (1200, 1000, 800, 600 bp), excluding positions with missing data and $USS_{10}$ that were 400 bp or less from positions with low input coverage (≤ 20 reads). Separation of ≥1000 bp was found to give the best compromise between good peak separation and the number of USS meeting the separation criterion (237 $USS_{10}$s and 209 $USS_{9.5}$s).

The search for non-USS sequences causing weak uptake effects used a subset of positions that were at least 0.6kb from the closest $USS_{10}$. This gave 575 'far from USS' segments summing to 29% of the genome. Uptake maps of segments containing positions with uptake ratios > 0.2 were examined visually to distinguish between (i) shoulders of adjacent $USS_{10}$ peaks, (ii) increased uptake at USS with scores between 9.5 and 10, and (iii) increased uptake at non-USS sequences.

**Incorporating within-USS interaction effects into uptake predictions:** Figure 6 of Mell et al. (Mell et al., 2012) shows the strength and direction of pairwise interaction effects between positions on the same USS , inferred from uptake analysis of synthetic degenerate USS. From this figure we extracted the mid-range value of the interaction effect at each interacting pair of USS positions (only some pairs of positions showed such effects). For each 86-028NP $USS_{9.5}$ whose sequence differed from the USS consensus at both positions of such a pair, the USS score was modified by adding or subtracting the corresponding interaction value. The modified scores were then used by the model to predict DNA uptake, as described above.

**Simulated noise analysis:** Simulated noise-free uptake data for short and long fragments was first generated by smoothing raw uptake coverage data for 86-028NP-short (sample UP7) and 86-028NP-long (sample UP3) using a LOESS regression, and normalizing the results to a mean coverage of 1.0. Simulated relative-noise amplitudes for every genome position were generated using the 'tuneR' R-package (Ligges et al., 2018). Before being added to the noise-free data, the noise amplitude at each position was adjusted in proportion to the noise-free simulated coverage at that position, with the maximum noise range for each coverage level set by a multiplier (1.0, 1.5, 2.0, 2.5 or 3.0) and by the range of all experimental replicates for positions with that mean coverage.  Red noise was used because, when added to the simulated noise-free coverage it gave an autocorrelation of 0.999, identical to that of the experimental data. Other noise types were evaluated but not used, since their autocorrelations were lower (0.975 for pink noise and 0.836 for white noise).
